# ChromatinHD connects single-cell DNA accessibility and conformation to gene expression through scale-adaptive machine learning

**DOI:** 10.1101/2023.07.21.549899

**Authors:** Wouter Saelens, Olga Pushkarev, Bart Deplancke

## Abstract

Machine learning methods that fully exploit the dual modality of single-cell RNA+ATAC-seq techniques are still lacking. Here, we developed ChromatinHD, a pair of models that uses the raw accessibility data, without peak-calling or windows, to predict gene expression and determine differentially accessible chromatin. We show how both models consistently outperform existing peak and window-based approaches, and find that this is due to a considerable amount of functional accessibility changes within and outside of putative cis-regulatory regions, both of which are uniquely captured by our models. Furthermore, ChromatinHD can delineate collaborating regions including their preferential genomic conformations that drive gene expression. Finally, our models also use changes in ATAC-seq fragment lengths to identify dense binding of transcription factors, a feature not captured by footprinting methods. Altogether, ChromatinHD, available at https://deplanckelab.github.io/ChromatinHD, is a suite of computational tools that enables a data-driven understanding of chromatin accessibility at various scales and how it relates to gene expression.

## 2. Introduction

Changes in DNA accessibility are a major hallmark of gene regulation [1, 2], and techniques that combine chromatin accessibility with RNA sequencing are opening up novel avenues to explore the interplay between transcription factor binding and chromatin state changes in influencing gene expression [3]. These methods not only facilitate the dissection of intricate gene regulatory networks [4], but they also illuminate the extent of intercellular and intracellular variability in transcriptional and epigenomic states [5, 6]. Furthermore, these techniques have great potential for both fine-mapping [7] and functional understanding [8] of non-coding genetic variation.

One of the very first steps of analyzing chromatin accessibility typically involves binning the raw data into putative cis-regulatory elements (CREs), using peak-calling [9–12], predefined regions [4, 13] or sliding windows with a predefined size [14–16]. This critical preprocessing step is based on the idea that evolution created distinct modular regions in the genome that are regulated in a coordinated way across cell types through transcription factor binding, chromatin modifications, nucleosome displacement and/or chromatin interaction changes [13]. The CRE, often referred to as an enhancer or promoter, is in this way seen as the fundamental functional unit of gene regulation, and it is thought that multiple such functional units often act in a combinatorial way to regulate gene expression [17]. Discretizing accessibility information into such putative CREs facilitates downstream data analyses because common statistical methods for differential expression, batch correction, dimensionality reduction, correlation analysis, differential transcription factor (TF) binding, and predictive modeling of gene expression can rapidly consume a CRE-based count matrix [11, 12, 18–22].

However, there is increasing evidence that reducing gene regulation to a coordinated effort of modular CREs may be an oversimplification. Multiple studies have highlighted inconsistencies and limitations of peak calling across cell types and methods [10, 11], both within and outside of canonical CREs [6, 13, 15, 16]. From an experimental perspective, there is an extensive body of evidence that gene regulation involves very different scales [23]: combinatorial TF binding localized at a few dozen base pairs [24, 25], nucleosomal interactions at 100bp scale [26], and genome organization [27, 28] combined with hub formation at kilobase scale or higher [29, 30]. These different scales indicate that *a priori* summarization at the CRE level, typically encompassing several hundred base pairs, may be too reductive to illuminate the full regulatory landscape underlying gene expression.

We developed ChromatinHD, a suite of computational methods that performs predictive and differential analysis of single-cell ATAC+RNA-seq data using the raw fragment data. Rather than making *a priori* assumptions about how the data should be structured, ChromatinHD uses neural network architectures and normalizing flows to automatically determine the functional regions and appropriate resolution to describe those regions in a cell type/state and position-specific manner. We apply ChromatinHD to show that there are inherent biases to current CRE-centric approaches, which affect the functional and mechanistic interpretation of chromatin data. Compared to these approaches, our scale-adaptive models are more predictive for gene expression and identify differentially accessible regions (DARs) that are more strongly enriched for functional binding sites and functional genetic variation. We include a series of interpretation tools that can be used to gather information about which regions are likely important in a given cell type, state or cell. Finally, we highlight how ChromatinHD captures information on (1) the juxtaposition between co-predictive enhancers and DNA contact, which are indicative of preferential genomic conformations driving gene expression, and (2) dense TF binding that is visible through changes in fragment sizes but not captured by typical footprinting approaches. Altogether, our data-driven approach provides the same interpretability as CRE methods, but extracts more fundamental gene regulatory information that would be missed otherwise.

## 3. Results

### 3.1 ChromatinHD enables scale-adaptive scATAC+RNA-seq analysis

By contemplating raw single-cell ATAC+RNA data, it is clear that the reorganization of open chromatin across cell states involves multiple genomic scales, with changes in gene expression and accessibility co-occurring within less than 100 base pairs (**Figure ED1a**), within 1kb (**Figure ED1b**) or across several kilobases (**Figure ED1c**). Furthermore, changes can occur asymmetrically within “peaks” that are present across various cell types (**Figure ED1c**), and, although this is harder to visualize, it may be that both fragment size information and co-occurence of fragments could provide valuable information about how these accessibility changes modulate gene expression. With this in mind, we designed two models (**Figure 1a-b**) that can capture these different features to predict gene expression or determine differentially accessible regions by using single-cell ATAC+RNA data as input.

In *ChromatinHD-pred*, we use raw chromatin accessibility fragments as input to predict gene expression (**Figure 1a**), which enables pinpointing which accessibility features, such as the position and fragment size, are predictive for gene expression. Here, we leveraged concepts from transformer models [31], which convert absolute positions of objects, typically text, into a positional encoding that can then be consumed more readily by downstream neural networks. In our case, these objects correspond to individual Tn5 cut sites relative to the canonical transcription start site (TSS) of a gene. ChromatinHD provides this positional encoding to a neural network that will learn which resolution is most relevant (**Figure 1c**), and pool information across different fragments from the same cell to predict gene expression (Methods). By looking at multiple fragments at the same time together with the corresponding gene expression, *ChromatinHD-pred* will automatically choose to focus on large or small regions (**Figure 1c**), located close or far away from the TSS, while also conditioning on the fragment size (**Figure 1d**), all in function of other fragments that are present in the same cell (**Figure 1e**).

In *ChromatinHD-diff*, we learn the difference in accessibility between different cell types, individuals, or conditions (**Figure 1b**), which can provide information about the activity of TFs [4]. Current tools typically do this by aggregating information across cells into “pseudobulk”, followed by a statistical model, e.g. a Wilcoxon rank-sum test, t-test, logistic regression or more complex generalized linear models [11, 20]. To make this approach scale-adaptive, we leveraged concepts from normalizing flows [32, 33], which are used to model complex probability distributions with a tractable likelihood that can be used for scalable statistical training and inference. In summary, *ChromatinHD-diff* will model the probability of finding a Tn5 insertion site using a series of reversible transforms that transform a simple uniform distribution into a complex distribution. Each of these transforms works at a different resolution (∼10kb, ∼1kb, ∼100bp), although these can be fine-tuned further by the model in a position-specific manner, and has a set of cell type/state-specific learnable parameters to allow small or large changes depending on the cell type/state (Methods). As such, *ChromatinHD-diff* not only models the distribution of insertion sites, but also how this distribution changes in different conditions, even if the changes in accessibility are discordant with respect to position, resolution, or across cell types (**Figure 1f**).

### 3.2 ChromatinHD outperforms current methods for analyzing scATAC+RNA-seq data

We first set out to comprehensively benchmark ChromatinHD models to the current state-of-the-art workflows, which involve defining CREs using peak calling, window scanning or predefined regions [12], followed by differential accessibility or predictive modeling using the cell-by-region count matrix.

First, we evaluated *ChromatinHD-pred*’s ability to predict single-cell gene expression using chromatin accessibility (**Figure 2a**), compared to CRE-based approaches that use peaks, windows or predefined regions. Direct comparison with different methods is challenging, because most methods, such as Signac [11], ArchR [20], or window-based approaches [15], do not directly use all peaks/windows to predict gene expression.

As discussed in-depth in the methods, we created a benchmarking set-up that most closely tries to recapitulate what these methods internally do, and further expanded to also include state-of-the-art regularized and non-linear regression to provide a comprehensive benchmark. We also included several baseline methods that count the number of fragments close to a gene’s TSS without prioritizing certain regions (methods). We found that ChromatinHD is the only method that consistently outperforms all other methods (**Figure 2b**) across most datasets (**Figure S1**). We note that because both the input (ATAC-seq counts) and output (normalized RNA-seq) are both very sparse and noisy, the performance differences are relatively small, with ∆*cor* around 0.01. Nevertheless, to ensure that this predictive performance was robust between technical and biological replicates, we tested performance on several datasets unseen by the training process, which yielded a similar performance if the same cell types were present (**Figure 2b**, middle). When transferring between datasets containing different related cell types/states, ChromatinHD’s performance reverted to the same predictive performance as the baseline, while still being higher than most other methods (**Figure 2b**, right).

**Figure 1:**
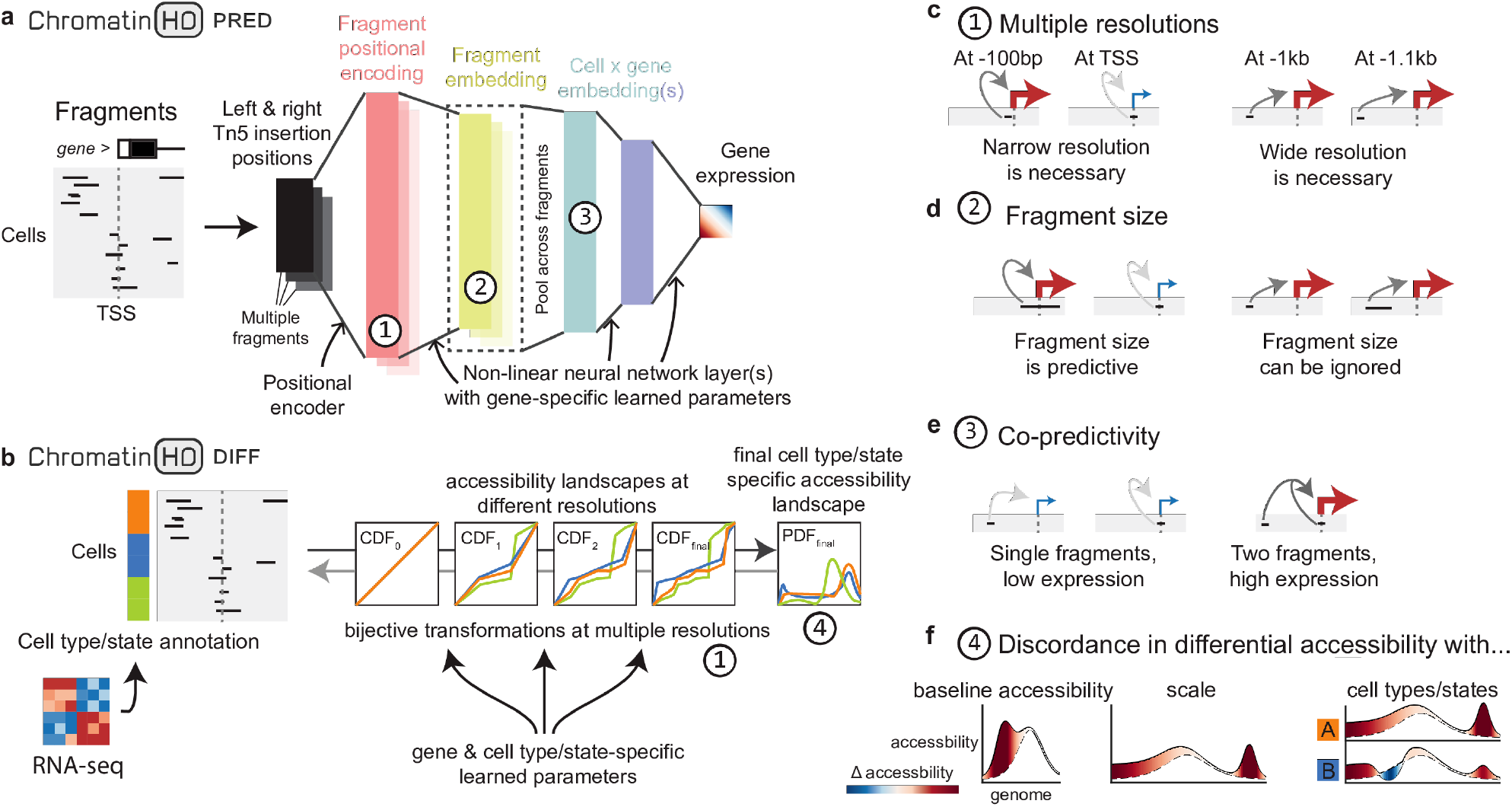
Central concepts behind *ChromatinHD-pred* and *ChromatinHD-diff* . (**a**) *ChromatinHD-pred* inputs raw fragments in a neural network architecture, that will (1) transform the positions of each fragment close to a TSS (e.g. -10kb or-100kb) into a positional encoding, (2) transforms this positional encoding into a fragment embedding, typically with a smaller number of features, using one or more non-linear neural network layers, (3) pools the fragment information for each cell and gene, (4) predicts the gene expression using one or more non-linear layers. Several features are highlighted by circled numbers as represented in *c-f*. (**b**) *ChromatinHD-diff* uses cell type/state annotation derived from, for example, single-cell RNA-seq to construct a complex multi-resolution cell type/state-specific probability distribution. To do this, we apply several bijective transforms on the cumulative density function (CDF), to ultimately be able to estimate the likelihood of observing a particular cut site using the probability density function (PDF). (**c**) *ChromatinHD-pred* can work at multiple resolutions, to capture changes within 100bp that do not (left) or do (right) affect gene expression. (**d**) *ChromatinHD-pred* can capture co-predictivity between two fragments within the same cell that may be predictive for gene expression in a non-linear way. (**e**) *ChromatinHD-pred* can also capture dependencies between fragment size and gene expression, for example to capture whether larger fragments are more predictive for gene expression than smaller fragments, in a position-specific manner. (**f**) *ChromatinHD-diff* can capture various complex differential probability distributions where differential accessibility could be discordant based on the position (left), resolution (middle) or cell type/state (right).

**Figure 2:**
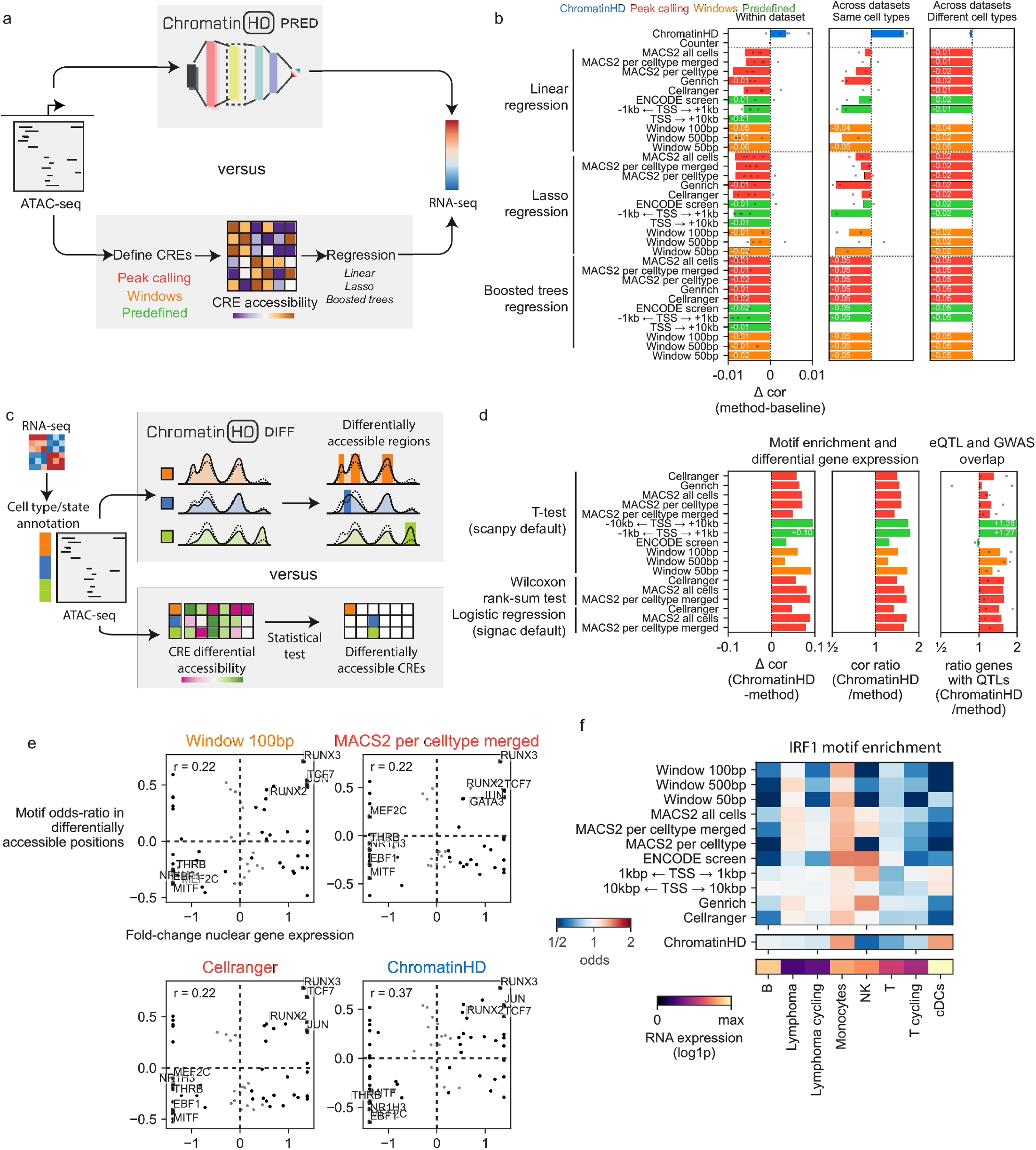
Benchmarking of *ChromatinHD-pred* and *ChromatinHD-diff*. (**a**) Evaluation design for predictive models. ChromatinHD is directly applied on raw fragments to predict gene expression (top). Peak-calling approaches, window scanning approaches with varying window sizes, or different pre-defined sets of cis-regulatory regions (CREs) are applied to construct a CRE accessibility matrix. The latter can then be used to predict gene expression using different regression methods (bottom). (**b**) Predictive accuracy (Δ*cor*) for different methods. Performance is relative to the baseline (counter), which consists of a simple linear regression between the total number of fragments and gene expression. (**c**) Evaluation design for differential models. ChromatinHD (top) directly calculates the probability density of cut sites, both the average across cell types/states (dotted line) as well as the difference (solid line), which are used to identify consecutive regions of accessibility. Other methods (bottom) typically first require a cell type/state by CRE matrix, after which a statistical test determines differentially accessible CREs. (**d**) We directly compared differential ChromatinHD regions with differentially accessible CREs, matching the former to comprise the same number of genomic positions as the latter. Differential models were evaluated on 3 metrics: the difference in correlation between motif enrichment odds and gene expression fold-change, the ratio of these two correlations, and the ratio of the percentage of genes where at least one eQTL/GWAS SNP was present in a differential CRE/region. (**e**) Enrichment of the IRF1 motif in various cell types and according to different methods that define differential accessibility. (**f**) Relationship between differential gene expression and motif enrichment in differentially accessible regions in T-cells from the lymphoma dataset.

Next, we evaluated how well different methods can capture differentially accessible regions (DARs) between two or more cell types or states (**Figure 2c**). As a proxy for a ground truth, we reasoned that differentially expressed TFs should be enriched in DARs, and thus formulated a score that correlates the change in gene expression with the enrichment/depletion in TF motifs. We found that ChromatinHD consistently outper-forms all other peak-, window- and predefined-based methods across datasets (**Figure 2d**). Averaged over all datasets and methods, ChromatinHD has about a 1.5x stronger correlation between differential motif enrichment and gene expression (**Figure 2d, Figure S2**).

We highlight several examples of improved TF binding site enrichment (**Figure 2e, Figure ED2a-b**). In some cases, the improved enrichment is simply quantitative, in the sense that its magnitude is stronger in ChromatinHD’s case even though all approaches predict the correct direction (**Figure 2e**). Still, we noted that the improved enrichment can be TF-dependent and that some TFs are not enriched in peak-based approaches, but are within ChromatinHD regions (**Figure ED2a**). This includes IRF1 in conventional dendritic cells in the lymphoma dataset, whose motif is not enriched or depleted according to peak/window based approaches, while ChromatinHD finds a strong enrichment that is concordant with its expression profile (**Figure 2f**) and literature [34]. This indicates that for some TFs, a peak-based approach may be sufficient to capture the relevant accessibility changes, while introducing biases towards those TFs that do not fit well within the peak paradigm.

We also evaluated DARs based on whether at least one immune-related GWAS SNP or expression-QTL was present in a DAR, and found that ChromatinHD captures on average 1.5x more GWAS SNPs and about 1.2x more eQTLs than other approaches (**Figure 2d, Figure S3**).

### 3.3 Interpreting ChromatinHD models highlight functional accessibility changes within and out-side canonical CREs

We implemented several interpretation tools (**Figure 3a-b**) for both ChromatinHD models that enable functional interrogation of the accessibility changes captured by the model. We first focused on interpreting genomic positions that are most predictive or differential, while interpretation of other aspects of the accessibility data (multiple fragments and fragment sizes) will be discussed later.

To interpret *ChromatinHD-pred* positionally, we censor the input data to remove fragments for which one of the Tn5 insertion sites fall within windows of various sizes, and then compare the test performance between the full and censored dataset (**Figure 3a**) (Methods). To interpret *ChromatinHD-diff* positionally, we calculated the marginal (differential) probability distribution across all factors except a particular window (**Figure 3b**) (Methods). By visualizing both positional predictivity (∆*cor*) and differential accessibility, we found that there is extensive functional heterogeneity within and outside CREs (**Figure 3, Figure ED3, Figure S4**). We cross-referenced with TF binding motifs, ChIP-seq data (whenever available) and immune GWAS SNPs, and often found a striking overlap between ChromatinHD regions and TF binding, both very localized (such as -7kb of the *CD74* TSS, **Figure 3c**), or within a broader region (such as the upstream gene body of *CD74*, **Figure 3c**).

**Figure 3:**
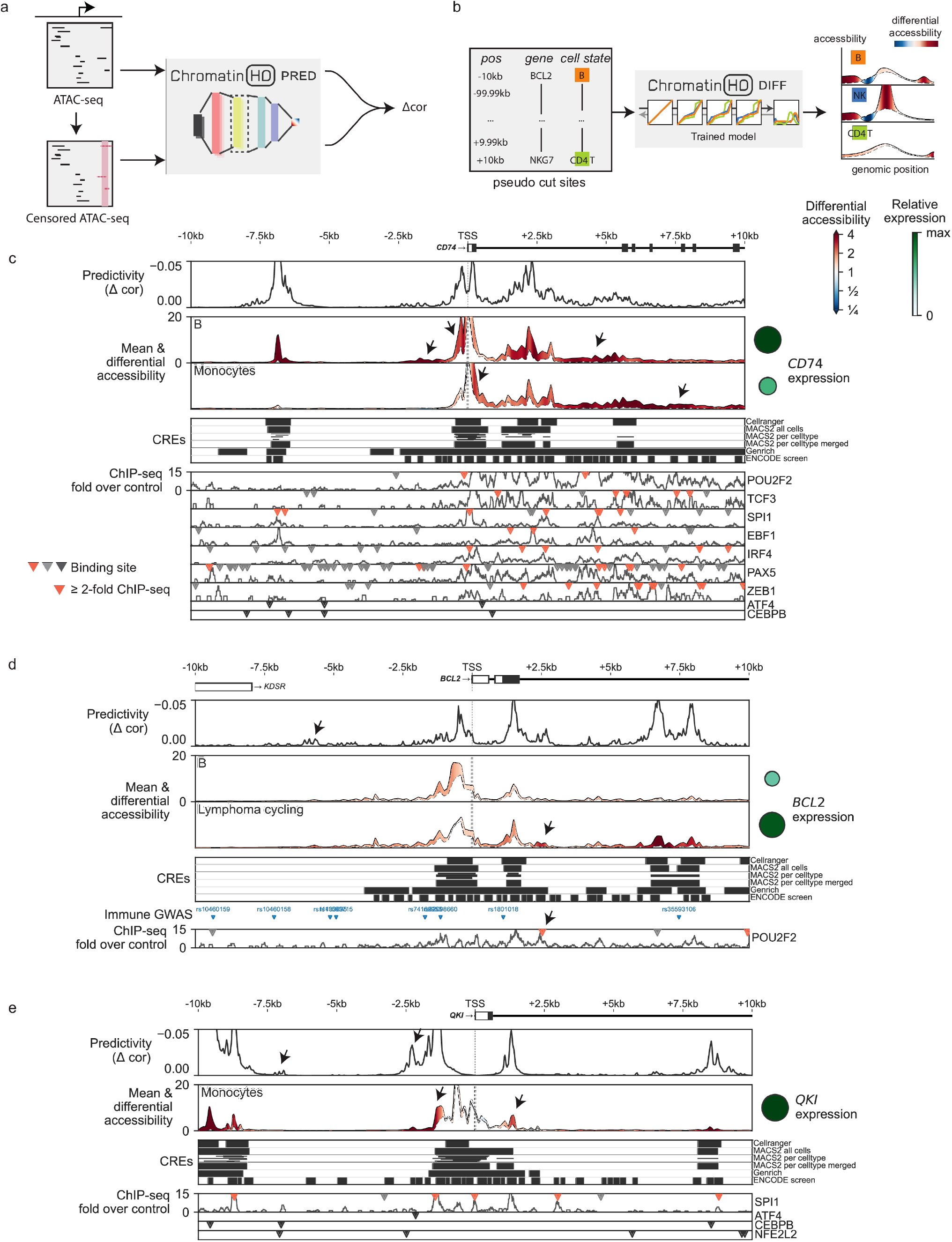
Interpretation of ChromatinHD models. (**a**) To interpret *ChromatinHD-pred*, we censor particular elements from the data and assess how it affects the predicted gene expression, i.e. the difference in correlation Δ*cor*. (**b**) To interpret *ChromatinHD-diff*, we create pseudo-cutsites at various positions, genes and cell states. We then use the trained model to calculate the probability of observing cut sites for each of these features, and compare these across cell types/states (vertically) and positions (horizontally). (**c-e**) Positional interpretation of both *ChromatinHD-pred* (Predictivity, top) and *ChromatinHD-diff* (Mean and differential accessibility, middle) models across different genes, along with positions of immune-related GWAS SNPs (colors representing different haplotypes with LD *r*^2^ *>* 0.9), motifs of relevant transcription factors and ChIP-seq tracks if available. (**c**) *CD74* has a strongly predictive and differential gene body, with numerous binding sites of transcription factors not captured within canonical CREs. Also note the asymmetry of predictive/differential accessibility around the BCL2 TSS between B-cells and Monocytes. (**d**) *BCL2* has a binding site of POU2F2 at +2.5kb from the *BCL2* TSS not captured by CRE approaches. (**e**) *QKI* has broad accessibility kilobases around the TSS, but only the outermost regions are predictive for gene expression, coinciding with those regions most strongly bound by SPI1 and, potentially, other Monocyte TFs. The arrows indicate the respective regions of interest.

Predictive and differential positions were frequently not included within any annotated CRE (**Figure 3c-e, Figure ED3a-c**), or were included within a larger CRE in which only a subset of positions were predictive/differential for gene expression in one or more cell types (**Figure 3e, Figure ED3a, Figure 3c**). We studied these “background” and “bystander” biases comprehensively, and found that they are present in all CRE-based methods (Supplementary Note 1). These biases are not simply theoretical, as they can have strong consequences on downstream interpretation. For example, IRF1 binding sites close to the *HLA-DQA1* TSS were strongly enriched within predictive regions and DARs, but only those within very accessible DNA were included within a CRE by most methods (**Figure S4a**), while ChromatinHD found several binding sites in predictive but less accessible regions. This exemplifies why *IRF1* is stronger enriched in differential Chromat-inHD regions (**Figure ED2a**). Another example is upregulation of the key anti-apoptotic factor BCL2 in B-cell lymphoma, which is known to be regulated by POU2F2, although its binding sites could not yet be established [35, 36]. Using ChromatinHD, we found a strong binding site of POU2F2 at +2.5kb from the *BCL2* TSS in clearly differentially accessible DNA, but which was not detected in any differentially accessible CREs either because the sites are eclipsed by nearby non-differential chromatin (MACS2 and Cellranger), or included in a very wide, mostly non-differentially accessible, region (Genrich) (**Figure 3d**). GWAS SNPs themselves were also frequently excluded from CREs even though the accessibility was clearly predictive and differential (+5kb *ZAP70*, **Figure ED3d**; -3.5 and -1.5kb *IGF2BP2*, **Figure S4b**). As a final example, the region around the *CD74* TSS is seemingly a single peak of about 1kb, but has two predictive elements, one upstream element primarily active in B-cells and bound by B-cell TFs (such as POU2F2 and PAX5), and a downstream element primarily active in Monocytes and containing motifs for Monocyte TFs (such as ATF4 and CEBPB) (**Figure 3c**).

Overall, these examples highlight that scale-adaptive methods such as ChromatinHD are important to fully capture the complexity of the regulatory landscape because CRE-centric methods frequently fail to detect intra- and inter-CRE heterogeneity. Despite involving a more complicated model, the interpretation of ChromatinHD models can reach the same level of interpretability as typical CRE-based methods, because, if need be, CREs can simply be defined after the model has been trained.

### 3.4 Co-predictivity identifies a 1-5kb outward juxtaposition between co-predictive regions and DNA contact

Because ChromatinHD’s predictive model uses a nonlinear multilayer mapping between one or more fragments and gene expression, it can capture information that goes beyond the position of individual cut sites, such as the presence of other fragments within the same cell (**Figure 1e**) or the size of the fragments (**Figure 1d**). We first assessed whether ChromatinHD can capture dependencies between fragments within the same cell, by comparing a model that can only additively share information across fragments with a model that can do so in a non-additive way. Even though the co-occurrence of multiple fragments close to a TSS is a rare occurrence (**Figure ED4a-b**) we found that the non-linear model performs better (∆ cor > 0.05, 13.4%) or approximately equal (0.05 > ∆ cor > -0.05, 86.6%) for the large majority of genes (**Figure 4a**). Genes that gained from a non-additive model were typically those that already had a high predictive performance with the additive model, indicating that the additive model reaches a saturation point which can be overcome by sharing information from multiple fragments (**Figure 4a**).

To determine whether two regions in the genome are co-predictive for gene expression in the same cells, we correlated the predictive accuracies between pairs of genomic windows (100bp) across cells, allowing us to identify “co-predictivity” as a measure of putative cooperation between regions (**Figure 4b-d**). We found that the majority of co-predictive pairs work synergistically, meaning that if fragments from two positions co-occur in a cell, they are typically in agreement on how the gene will be expressed (**Figure ED4c**). Still, negative co-predictivity does occur in a significant set of cases, and is generally associated with regions where changes in accessibility and gene expression are inversely correlated (**Figure ED4c**), indicative of differential usage of enhancers between cell types/states. We found that positive co-predictivity is higher within shorter genomic distances (<20kb) (**Figure 4e**) and that the region around the TSS has a slightly higher average co-predictivity compared to up-or downstream regions (**Figure 4f**). Genes that gained performance in the non-additive model typically had strong co-predictivity with the TSS (**Figure ED4d**, e.g. *KLF12* and *TNFAIP2*, **Figure 4b-c**) and co-predictivity was more associated than overall predictivity with the presence of GWAS SNPs in one of the two regions (**Figure 4g**), suggesting that these pairs of regions have a biomedical relevance. Co-predictivity can also inform on how seemingly distinct genetic variation can converge through a similar mechanism. For example, the first *BCL2* intron contains 3 independent SNPs not under linkage disequilibrium in distinct co-predictive regions, all associated with granulocyte numbers [37–39], suggesting that these distinct genetic variants work through a common regulatory hub to regulate downstream gene expression (**Figure 4d**).

With co-predictivity, we have a readout on cooperativity between regions that is conceptually distinct from both DNA contact frequencies [27, 40–42], and DNA proximity [43]. Several recent studies have shown how these latter methods can produce seemingly paradoxical results, with the physical proximity between specific enhancer and promoter pairs decreasing upon active transcription, while DNA contact frequencies increase [44–46]. This decreased proximity may be related to the establishment of a high protein concentration environment [47]. However, whether this is a general genome-wide feature of *in vivo* mammalian gene regulation is not well understood.

The high-resolution, genome-wide readout provided by co-predictivity may help reconciling these seemingly non-intuitive observations. We compared ChromatinHD’s co-predictivity matrices from the pbmc10k dataset with Hi-C data at 1kb resolution [27] originating from GM12878, a B-cell derived cell line. When investigating the Hi-C signal of individual co-predictive pairs, we found that these are often close but not exactly overlapping with physically contacting regions, frequently missing each other by 1 or several kilobases (**Figure ED4e**). Indeed, if we compared co-predictivity with max-pooled Hi-C signal, we found that the overlap was the highest at about 2-5kb of max-pooling (**Figure 4h**). This confirms that DNA contact is more frequent than random in (co-)predictive regions (of ∼1 kb), but that it is much higher in the adjacent regions, potentially to accommodate protein and RNA complexes required for gene regulation.

We next studied whether the juxtaposition of co-predictivity and DNA contact depends on their orientation, because this could help disambiguate different 3D conformations, and therefore help us understand how this interaction is established or maintained (**Figure 4i-q**). Assuming a given genomic distance between a co-predictive region pair (initially 20kb-25kb), a DNA contact at respectively -2kb and +2kb of the two co-predictive regions would be indicative of a loop which exposes the two co-predictive regions outward (**Figure 4j**). Indeed, after normalizing with random non-co-predictive regions, we found that the DNA contact is the strongest for such a configuration (**Figure 4i-j**), compared to the less frequent - but still enriched - contacts overlapping with the co-predictive regions themselves (**Figure 4k**), juxtaposed in the same orientation (**Figure 4l**) or juxtaposed to inside of the loop (**Figure 4m**).

**Figure 4:**
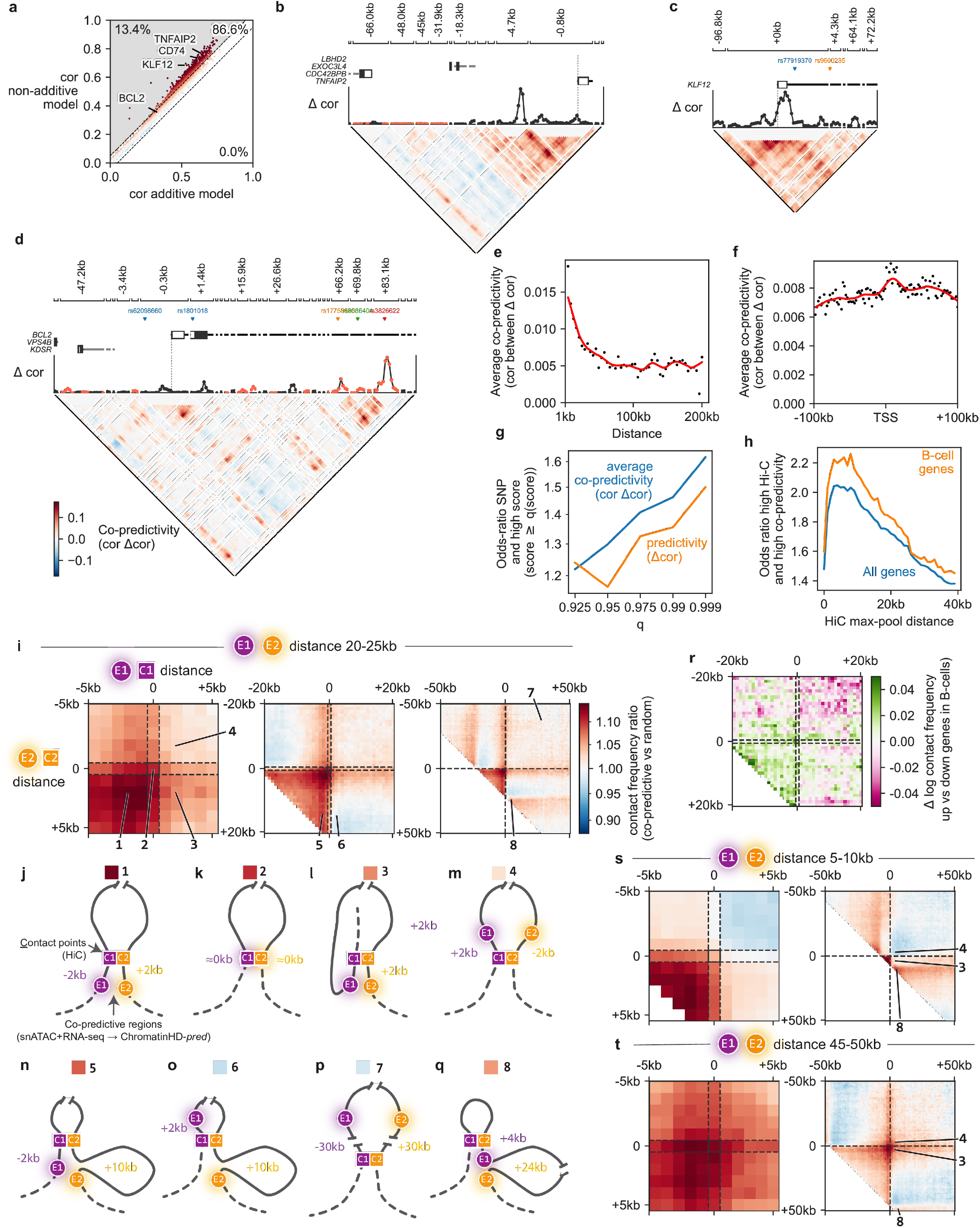
A 1-5kb shift in co-predictivity and DNA contact highlights preferential DNA conformations connecting two co-predictive regions. (**a**) Predictive test-set accuracy of additive and non-additive *ChromatinHD-pred* models across all genes. Percentage of genes with Δ*cor* higher or lower than 0.05 are indicated. (**b-d**) Examples of co-predictive regions for the *TNFAIP2, KLF12, BCL2* genes. Immune-related GWAS SNPs are shown and colored according to haplotype (LD *r*^2^ ≥ 0.9). (**e**) Average co-predictivity, i.e. the mean *cor*Δ*cor* across different 1kb bins. (**f**) Average co-predictivity for various distances between positions. Note that positions closer than 1kb were excluded. (**g**) Average odds-ratio across genes for finding an immune-related GWAS SNP in a top *q*% predictive or top *q*% co-predictive region. (**h**) We calculated for each gene and for each slice of genomic distances (1kb-10kb, 10kb-20kb, …), the odds-ratio for finding high co-predictivity (higher than median) and high Hi-C signal (higher than median). This was performed both for the original Hi-C data (Hi-C max-pool distance = 0), but also for max-pooled Hi-C data where we took the maximal Hi-C signal at various genomic manhattan distances around the original position. B-cell genes were defined as being differentially expressed in naive, memory or plasma cells compared to all other cell types in the dataset. (**i**) Hi-C pile-ups of potential DNA contact points (C1 and C2) close to two co-predictive regions (E1 and E2, distance 20-25b, *cor*Δ*cor >* 0). Shown is the relative Hi-C signal centered on the co-predictive pair divided by a random pair around the same gene with the same genomic distance. (**j-q**) Illustrations of how different distances between predictive regions and DNA contact points from *a-c* may relate to DNA conformation. (**r**) Difference in log contact frequency between up and down-regulated genes in B-cells. (**s-t**) Same as *a-c* but with E1-E2 distances of 5-10kb and 45-50kb respectively.

When zooming out to assess larger enhancer-contact point distances (**Figure 4i** middle and right), we found a preference for the co-predictive regions to be located outside of the contact points (**Figure 4n**), with DNA contact being less likely than random if they would result in the two regions being on opposite sides (**Figure 4o**), or deep (>20kb) on the inside of the loop (**Figure 4p**). Interestingly, we also saw an enrichment for situations where only one of the regions is close to a contact point as long as both co-predictive regions are located on the same side of the loop, indicative of further looping and hub formation (**Figure 4q**) [48, 49]. The dependency between co-predictive regions and DNA contact points changed depending on whether the regions were active in a cell, as co-predictive regions regulating B-cell-specific genes showed a more outwards dependency patterns in the B-cell-like cell line (i.e. **Figure 4j**), compared to the inwards pattern (i.e. **Figure 4m**) for those regulating genes expressed in other leukocytes (**Figure 4r**).

The general pattern of DNA contact between co-predictive regions was consistent across different sets of distances (**Figure 4s-t, Figure ED4f-g**) and was further validated on a recently published Hi-C dataset with higher resolution [41] (**Figure ED4h-i**). Notably, the inward pattern (**Figure 4m**) was disfavored for shorter distances (<15kb) (**Figure 4s, Figure ED4f**), while this pattern was more neutral or enriched with higher (>25kb) distances (**Figure 4t, Figure ED4g**), potentially because the larger stretch of DNA allows more physical freedom regarding how co-predictive regions can interact.

Altogether, by connecting co-predictivity analysis with Hi-C data, we have shown how co-predictive regions make contact in a slightly juxtaposed way, and that this juxtaposition is oriented in such a way to prefer looping and further hub formation. While this confirms and reconciles some of the results observed at individual loci [44–46], it also shows how ChromatinHD’s co-predictivity data provides a view on cooperativity between DNA that is complementary to both DNA contact and DNA proximity analyses.

The mechanism by which enhancer-enhancer and enhancer-promoter interactions are created and maintained is still controversial [50]. First, we found contacts in the same orientation are consistently enriched even over longer distances (**Figure 4i** right), indicating that some contacts between co-predictive regions may form independent of looping extrusion (**Figure 4l**). When contrasting highly and weakly co-predictive regions, we found that binding of looping factors RAD21, CTCF and YY1 are not enriched directly within highly co-predictive regions, but are enriched close (RAD21, CTCF, 500bp) or farther away (YY1, 1.5kb) to the co-predictive region, contrasting with the expected bell-shaped pattern of cell type-specific TFs (**Figure ED4j**). For example, YY1, described as a universal structural regulator of enhancer-promoter interactions [51], was depleted directly within the co-predictive regions (**Figure ED4j**). This further confirms the juxtaposition between co-predictivity and DNA contact, and that several mechanisms, CTCF-cohesin, YY1 and cohesin-independent, may all be at play to form or stabilize these interactions.

### 3.5 Large submononucleosomal fragments are indicative of dense TF binding and active gene regulation

Differences in fragment size is an additional feature that ChromatinHD enables us to consider (**Figure 1d**). This could be relevant since such differences have already been linked to distinct chromatin states and nucleosome positioning [52, 53]. To test this, we censored fragments of a particular length and assessed the effect of the model’s ability to predict gene expression. We found that while the predictivity of a fragment size is correlated to the number of fragments of that size in the data, there are clear relative differences in different “waves” corresponding to nucleosomal or sub/super-nucleosomal fragment sizes (**Figure 5a**). Averaged over all genes, nucleosomal fragments (160-190bp) were about 3 times less predictive than submononucleosomal fragments (80-120bp) (**Figure 5a-b**). We note that despite these relative differences, nucleosomal fragment sizes still contributed significantly to a model’s predictive performance and every fragment size was, on average, still positively correlated with gene expression at similar levels (**Figure 5b**).

Surprisingly, we saw a clear split in predictivity within nucleosome-free fragments, with TF footprint fragments (10-60bp) being much less predictive than larger submononucleosomal fragments (60-120bp, Mono−), despite the former being most frequent (**Figure 5a**) and typically strong indicators of TF binding [52, 54, 55]. We hypothesized that this may be due to the fact that TF footprint fragments mainly straddle isolated binding events on the genome, whereas the most relevant elements for gene regulation are at locations with concentrated direct and indirect TF binding, larger DNA protection, and therefore fragments that straddle longer regions [23]. To assess this, we contrasted motifs enriched in predictive regions to motifs enriched in regions with a high submononucleosomal versus TF footprint ratio. We found that nearly all TFs enriched in predictive regions are also enriched in high submononucleosomal regions, most of which are common regulators of specific immune cell types (**Figure 5c**). Only CTCF, a critical chromosomal organization regulator and the most common TF used in footprinting analyses [55, 56], showed a clear preference towards TF footprint fragments. To find further experimental evidence, we cross-referenced with TF footprints inferred using HINT-ATAC [54] on the same ATAC-seq data, and TF ChIP-seq data from GM12878, a B-cell-like cell line [13]. We found the number of TFs that bind (in)directly within a 100 bp window to be strongly positively correlated with both the predictivity of a window and the ratio of submononucleosomal versus TF footprint fragments, increasing linearly even for regions with dozens of bound TFs (**Figure 5d**). This was true both when we focused on TFs enriched in B-cells, as to best match the ChIP-seq cell line (GM12878) with the primary cells under study (PBMCs) (**Figure 5d**), but also when we considered all TFs (**Figure ED5a**). This increase in predictivity coincided with a decrease in the number of fragments, which further confirms the ambiguous relationship between accessibility and gene expression (**Figure 3**, Supplementary Note 1). Indeed, low accessibility can both mean high nucleosome occupancy (and therefore low expression) or high TF occupancy (and therefore typically high expression). By also considering fragment size in its model, *ChromatinHD-pred* is able to differentiate between the two cases that would seemingly look similar when one only considers fragment counts.

The use of footprinting methods has been controversial, particularly given several observations of functional binding events that do not leave footprints [57]. Our analysis highlights another challenge, namely that dense (indirect) binding of TFs may mask the local footprint signal, despite these densely bound regions being the most predictive for gene expression. Indeed, we found that the number of detected footprints stagnates and even decreases with an increasing number of TFs that bind to the respective regions (**Figure 5d, Figure ED5a**). We further verified this on bulk DNase I hypersensitivity datasets [55], which typically have a much higher sensitivity to detect footprints, confirming that these suffer from the same bias (**Figure ED5b**). Altogether, this shows that while footprinting methods can detect individual TF binding events, they are less able to distinguish densely bound regions from weakly bound ones, and in fact tend to be negatively biased towards the former. In contrast, although ChromatinHD does not provide direct evidence of TF binding, it is better able to detect these densely bound regions because the subtle shift in fragment sizes makes the model more predictive for gene expression.

**Figure 5:**
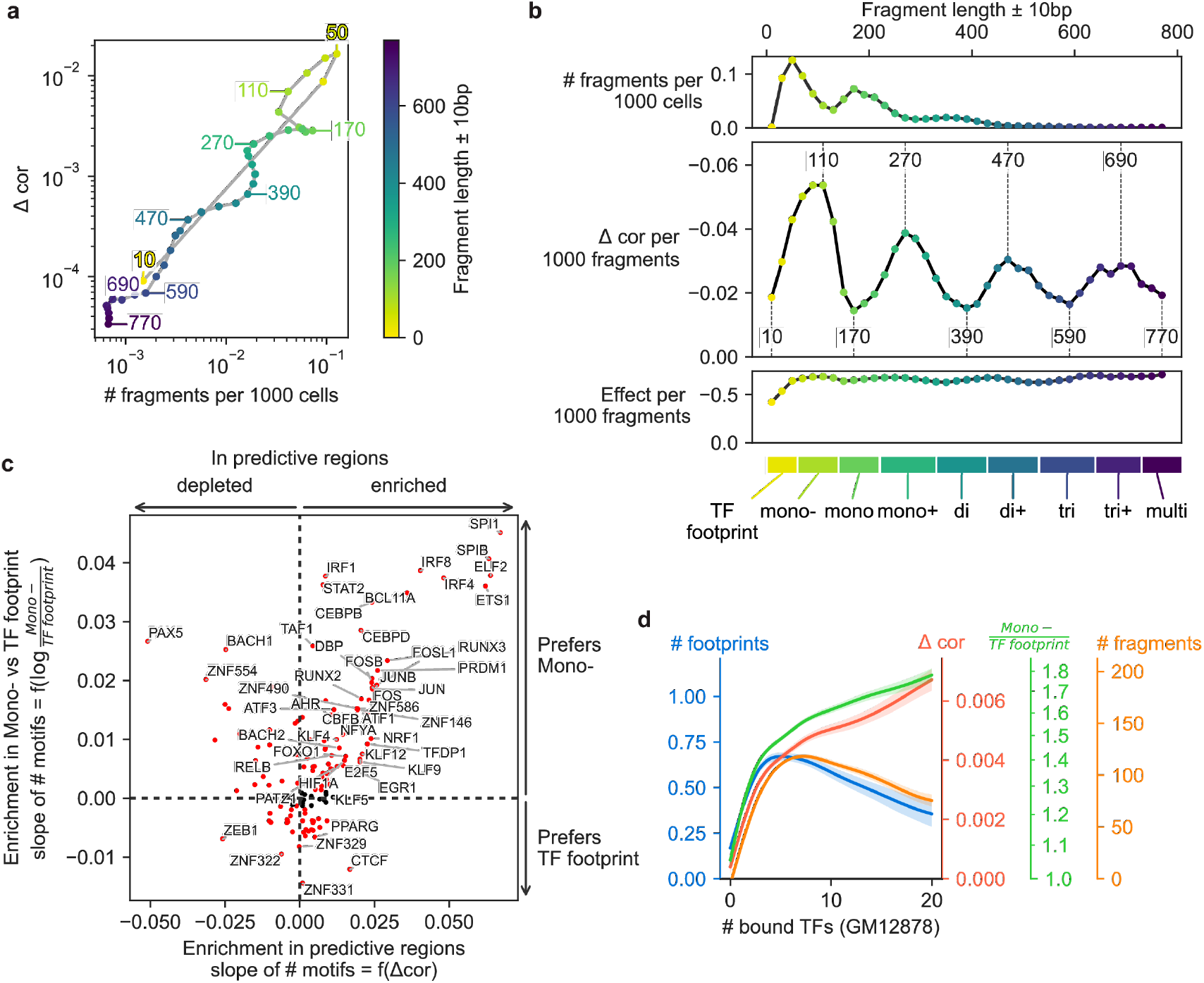
ChromatinHD learned a complex dependency between predictivity and fragment size. (**a**) Relationship between the abundance of a fragment size bin (± 10kb) and the overall loss in predictive accuracy (predictivity, Δ*cor*) when fragments of these sizes were removed from the data. (**b**) Abundance, normalized predictivity (predictivity divided by abundance) and average effect of different fragment size bins. On the bottom, a subdivision into different classes of fragments based on taking the middle point between local maxima and minima of normalized predictivity. (**c**) Motif enrichment for windows with Mono-(80-120bp) versus TF footprint (0-80bp) fragments. We first created a spline model that models the relationship between GC content and # of motifs within a 100bp window. Next, we modeled the relationship between the residual # of motifs and the log-ratio of Mono- and TF footprint fragments (x-axis). A similar procedure was applied to model the relationship between co-predictivity and GC-corrected motif frequency (y-axis). (**d**) Relationship of the # of (indirectly) bound TFs within a 100 bp window according to ENCODE GM12878 data and the # of footprints according to HINT-ATAC on the pbmc10k data (blue), predictivity (red), ratio of Mono-versus TF footprint fragments (green) and overall number of fragments (orange). Shown is the mean and standard error of a spline fit using R’s gam function with smoothing parameter *sp* = 1. ChIP-seq data of top 30 TFs was used, ordered by the correlation between predictivity and number of binding sites within 100 bp windows; data for all TFs is shown in **Figure ED5a**

## 4. Discussion

A central challenge in understanding eukaryotic gene regulation is learning how various chromatin state scales are integrated to regulate a gene’s expression [58]. An important advance toward addressing this challenge is the development of multi-omic profiling assays which directly link gene expression to genomic read-outs in the same cell [3] as these can inform both on TF binding and long-range CRE interactions [12]. However, his potential has so far mainly been exploited with methods using CREs as the main preprocessing step, which is likely too reductive to capture the full complexity underlying gene regulation. To address this, we have developed two machine learning models and interpretation tools that can contribute to learning how chromatin accessibility relates to gene expression.

By staying close to the actual biochemistry, i.e. raw fragments, we have shown that accessibility data contains information that goes well beyond changes within and outside canonical CREs. Although we do find that predictive changes in accessibility are indeed restricted to specific regions in the genome, our study mainly calls into question whether *a priori* defined CREs can be used as a summarization method to comprehensively study gene regulation [13]. The high probability that both “bystander” regions are included and functional “background” regions are excluded, means that summarization at the CRE level induces an undesirable bias that can affect the prioritization of regions for gene regulation, the identification of relevant TF binding sites and the fine mapping of genetic variants. On the other hand, by learning which regions are predictive and differential in a data-driven and scale-adaptive way, ChromatinHD models can better delineate these functional regions.

Furthermore, the high resolution offered by both ChromatinHD and deep Hi-C data, also allowed us to identify a juxtaposition of 1-5 kb between DNA contact and enhancer activity, which is likely related to preferential chromatin conformations that lead to gene regulation. Our data is consistent with previous observations made for individual loci [44–46], and suggests that these juxtapositions occur across the whole genome in vivo in mammalian DNA. *ChromatinHD-pred* also learned a strong predictive preference for longer sub-mononucleosomal fragments to predict gene expression, a chromatin accessibility feature that is an indicator of a very active regulatory environment with potentially high (in)direct protein binding, but which are less captured by footprinting methods. Both of these examples highlight how an unbiased, data-driven approach is useful to uncover additional, often hidden, layers of the complexity underlying gene regulation. As such, our study joins several other recent papers that have shown similar potential using window-based approaches [16], and footprinting analysis [6].

Active transcription often leads to opening of the gene body (**Figure 3**) and a limitation of ChromatinHD may be that it identifies regions that are not necessarily causal for gene regulation, but a consequence of active transcription. Still, whether this gene body accessibility increase is purely consequential is unclear, especially since we can observe strong ChIP-seq signal in these regions overlapping with putative binding sites (**Figure 3**), and several studies have recently showcased the role of RNA polymerase in enhancer-promoter interactions [59] and RNA-TF-mediated gene expression control [60]. In the future it may be beneficial to leverage perturbational data, e.g. utilizing multiplexed CRISPR [61] or natural variation [62] to help the model with distinguishing causal from consequential signals.

To enable the community to use and extend ChromatinHD, we made the PyTorch models, training and interpretation tools available as a python package (https://github.com/DeplanckeLab/ChromatinHD), making them easily deployable and extensible. With this package, users will be able to train, infer and interpret the models. For downstream analyses, e.g. gene regulatory network inference [4] or velocity analysis [5], users can directly plug the identified predictive regions in these tools. Finally, there is a great potential to expand ChromatinHD models to include dynamics [3], multiomics velocity [5], impact of genetic variation [17] as well as integrating spatial information [63].

## 5 Methods

### 5.1 ChromatinHD

ChromatinHD is available as a Python package at https://github.com/DeplanckeLab/ChromatinHD.

### 5.2 Data preprocessing

All six multiome datasets, (pbmc10k, pbmc10_gran, lymphoma, brain and e18brain, pbmc3k) were preprocessed in the same manner. Raw fragments, mapped either to the GRCh38 or mm10 genomes, and the raw expression counts at the gene-level, were obtained from the 10X Genomics website (https://www.10xgenomics.com/resources/datasets). Cells were filtered on containing at least 1000 UMIs and 200 genes with at least 1 UMI. We selected the 5000 most variable genes for downstream analysis by ordering on the normalized dispersion calculated by scanpy [64]. We obtained the canonical transcription start site (TSS) for each gene from biomart [65]. From this TSS, we extended either 10kb or 100kb up- and downstream to define a gene region. We then converted fragment location information in an efficient sparse data format with easy access to fragments from a particular minibatch of cells, and to which cell, genes or cell-by-gene combinations these fragments belong. This data format contains index pointers [66], indicating the first index at which the fragments from a particular cell begin.

### 5.3 ChromatinHD-pred

As input, *ChromatinHD-pred* (**Figure 1a**) uses a matrix containing the start and end positions of the fragments *X* and a mapping *M* for each fragment indicating to which cell and gene (locus) each fragment belongs. Fragments can be contained multiple times in this data format if they overlap multiple gene loci. The goal of *ChromatinHD-pred* models is to predict gene expression by focussing on fragments in a small or large region located around a predefined window relative to the TSS. While a network could in theory learn this from the raw positions, ChromatinHD uses a positional encoding, as used in sequence models [67], to more efficiently present the positions to the downstream neural networks. This positional encoding will convert an integer position of start and end Tn5 insertion sites into a set of continuous features. This encoder allows downstream linear and activation functions to easily learn to prioritize certain positions. We used the sinusoid positional encoding as used by the majority of modern transformer models [31]. For a position of a Tn5 insertions site *x*, we define the different dimensions *j* of the encoding as:

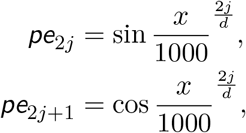

where *d* denotes the number of positional encodings and is typically chosen high enough so that a whole range of small and wide resolutions are available for the downstream neural network to choose from, in our case *d* = 50.

Next, within the model, the set of positional encodings for all fragments with dimension 2*d* is presented to a fragment embedder, which represents a set of neural network layers with gene-specific parameters that learns which positions of Tn5 cut sites may be relevant for a particular gene’s expression. For all models in the manuscript, we used two feedforward layers with 10 hidden dimensions. Next, the model pools information for all fragments into a cellxgene embedding by summing the fragment embedding for each cell and gene. If no fragments were present, the cellxgene embedding simply outputs zeros. This cellxgene embedding is then presented to another feedforward network, again with 10 hidden dimensions, that ultimately predicts gene expression.

To train the model, we split the data into train, validation and test cells according to a 3:1:1 split. We trained using minibatches of cells (n=100), and calculated for each epoch the correlation between predicted and observed gene expression on the validation set for each gene. Once validation performance increased relative to the previous epoch, we performed early stopping of the training for that particular gene. As loss function, we used the negative correlation between the actual gene expression versus the predicted gene expression. For the benchmark, we used normalized data, as this is the typical data used by archR [20] and signac [11]. For all downstream results, we used MAGIC imputed data [68] as implemented in scanpy [64] because this led to more stable results across different folds. We trained the model multiple times on 25 folds, constituting 5 different cell permutations. Parameters were optimized using a custom sparse ADAM [69] implementation, with learning rate 10^−3^ which ensures the update of gene-specific parameters only if fragments of a gene were observed during the forward pass.

### 5.4 ChromatinHD-diff

To learn how the accessibility changes between different cell types/states we defined a likelihood for observing a Tn5 insertion at position *x* in gene *g* in cell type *ct* as follows:

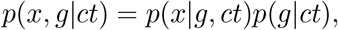

where *p*(*g*|*ct*) ∼ Categorical(*ρ*_*g*,ct_) captures the total increase or decrease in number of fragments between the cell types and genes. To parameterize this distribution, we first calculated 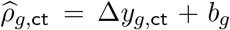 where ∆*y*_*g*,ct_ ∈ ℝ was optimized as a free parameter and regularized by using hierarchical Bayes with prior ∆*h*_ct_ ∼ Normal(0, 1). *b*_*g*_ was fixed to the average number of fragments present in a gene region over all cells. 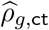 was transformed into probabilities using a softmax transform: 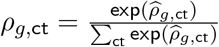

Determining *P* (*x*|*g*, ct) requires a multimodal density function, with numerous local minima and local maxima pertaining to broad or narrow areas of TF or nucleosome positioning (**Figure 1f**). For this, we use concepts from normalizing flows [32, 33] which use a set of bijective transformations on the cumulative density function to transform a simple distribution into a complex multimodal one. Specifically, we used three quadratic spline transforms of 32, 64, and 128 knots, where each transform is able to capture small (∼ kilo-base), medium (∼ several hundreds of bases) and small (∼ 100 bp or less) changes in accessibility, within a window of 20 kilobases, respectively.

Quadratic splines were parameterized by a set of unnormalized heights 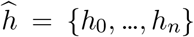 and widths 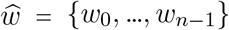where *n* corresponds to a predefined number of knots. Widths and heights were normalized as

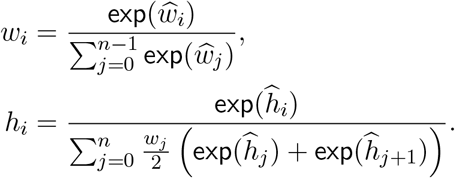

This normalization is necessary to maintain that the domain of the monotonic quadratic spline remains ∈ [0, 1], which ensures that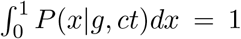 Next, we defined the locations for each bin as the sum of weights in the range if 1 to *i*, where *i* corresponds to the bin index and the left cumulative density function (CDF) for each bin as lcdf 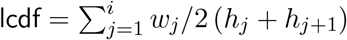

To calculate the likelihood of observing a cut site *P* (*x*|*g, ct*), we iteratively apply the quadratic spline transformation to both position *x*_*k*_ and its (log-)probability *p*_*k*_, with *k* ∈ {0, 1, …, *m*} and *m* being the number of transformations, in our case *m* = 3. Starting from *p*_0_ = 1 and *x*_0_ = *x*, we defined the coefficients of quadratic function applied for each cut site as:

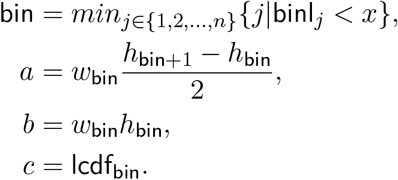

The transformed position and probability density can then be calculated by first calculating the normalized position of *x*_*k*−1_ as

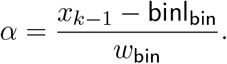

The transformation can then finally be applied as

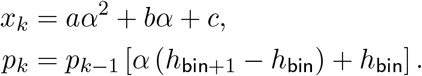

While the widths of each bin are shared among all cells, the unnormalized height 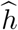 depends on the cell type-/state of the cell as 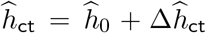 where 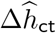 ∈ ℝ were free parameters learned during training and regularized by using hierarchical Bayes with prior 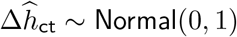.

To train the model, we split the data into train, validation and test cells according to a 3:1:1 split. We trained using minibatches of cells (n=100). As loss function, we used the evidence lower-bound:

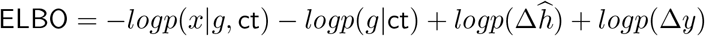

### 5.5 Interpretation of *ChromatinHD-pred*

To interpret *ChromatinHD-pred* models, we compared the test-set on the full data with that obtained from censored data. For positional predictivity in particular, we censored fragments for which a fragment was removed if one Tn5 insertion site was overlapping with a window. We performed this censoring using a scanning window approach of window sizes 50, 100, 200, 500, 1000 and 2000 base pairs, with a stride length of half of the window’s size. We calculated the robustness of a window’s ∆*cor* using a one-sided t-test across the different folds, and set the ∆*cor* to 0 if the adjusted p-values (Benjamini Hochberg correction) were higher than 0.1. We then extracted a base-pair position importance by linearly interpolating the ∆*cor* for a given window size, and summing up these interpolated values divided by the window size.

For interpretation of fragment size importance, we removed fragments of a particular size window (20 bp) ranging from 10 to 770, and compared the fragment size mean ∆*cor* across genes with the mean number of fragments of that particular size to a relative importance *rel*∆*cor*. To split into different types of fragments, e.g. TF footprints and Mono−, we calculated the local minima and maxima of *rel*∆*cor*, and split the fragment sizes by taking the midpoint between these minima and maxima.

### 5.6 Interpretation of *ChromatinHD-diff*

To interpret *ChromatinHD-diff* models, we use the trained model to extract *P* (*x, g*|ct) for all genes *g*, cell types ct and positions (with step size 25 bp). From this, we obtain a position-specific fold-change by calculating

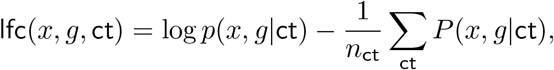

and linearly interpolating across positions. Differentially accessible regions can then be called by determining consecutive positions where lfc remains above a particular cutoff, by default log 2. When directly comparing differentially accessible CREs with differentially accessible ChromatinHD regions, this cutoff was adapted on a per cell type basis such that the number of differential ChromatinHD regions was the same as the number of differential CRE positions. We only retained regions where baseline accessibility was higher than a specified cutoff, i.e. *p*(*x, g*|ct) ≥ 1.

### 5.7 Benchmark

#### CRE methods

We included the most commonly used peak-calling approaches for single-cell RNA-seq data, i.e. MACS2 [70, 71], MACS2 merged across different cell types [11], Cellranger [72], and Genrich [73]. We also included SCREEN regions [13] as these have been proposed to be a good substitute for dataset-specific peak calling [4, 13].

For “MACS2 all cells”, we used MACS2 v2.2.7.1 [70], with fragments and the ‘–no-model’ and ‘–BEDPE’ parameter. For “MACS2 per cell type” we split the fragments based on the cell type, called peaks, and kept all peaks even if they overlap as separate features for downstream analyses. For “MACS2 per cell type merged”, the typical recommended pipeline [11], we merged all peaks from “MACS2 per cell type” using the bedtools merge [74]. For Cellranger, we used the peaks as provided by Cellranger 3.0.2 multiome pipeline. We used Genrich v0.6.1 with ATAC-seq mode ‘-j‘. For window approaches, we simply divided the region around a gene’s TSS in equal parts based on the window size. ENCODE SCREEN v3 data was obtained from https://screen.encodeproject.org/. Cell by CRE counts were obtained by determining whether one or two cut sites of a fragment overlapped with the CRE.

We used the CRE counts overlapping with (part of) a gene’s window to perform linear and lasso regression using scikit-learn’s ‘LinearRegression’ and ‘LassoCV’ respectively. For LassoCV, we provided the same set of validation cells as provided to *ChromatinHD-pred*, and ‘n_alphas=10‘, which will automatically determine the optimal penalization parameter on a per-gene basis. For boosted trees regression, we used XG-Boost’s ‘XGBRegressor’ with 100 estimators and 50 early stopping rounds. To perform this early stopping, we provided the same validation cells as provided to *ChromatinHD-pred*. Final correlation values were then obtained for test cells. If no CREs were found within the gene region, the correlation was set to 0.

All methods were run using 5-fold cross validation across the cells, with a 3:1:1 split for train, validation and test cells. For training and testing on different datasets, we applied training and testing on respectively the full train and test dataset.

For differential accessibility, we provided the CRE counts for those CREs overlapping with at least one gene window to scanpy v1.9.1 and signac Signac v1.9. We calculated differentially accessible CREs using (1) scanpy’s ‘rank_genes_groups’ for t-test, (2) ‘rank_genes_groups’ with ‘method=wilcoxon’ for wilcoxon test, (3) Signac’s ‘FindMarkers’ with ‘test.use=‘LR’’ and ‘latent.vars=‘peak_region_fragments’’ as specified in the tutorial (https://stuartlab.org/signac/articles/pbmc_vignette.html#find-differentially-accessible-peaks-between-clusters).

#### Comparing gene expression prediction

To make the comparison fair for all methods, we included all CREs that overlap at least partially with a given window around a TSS, in this case -10 kb and +10kb. Peak-based approaches are typically linked to target gene expression on an individual basis using correlation analyses [11, 20], and to best replicate this we included both linear and non-linear boosted trees regression. For window-based approaches both single-window correlational analysis at about 50-100 bp [15] as well as multiple window regularized regression using 500 bp [16] have been described, and as such both were included within the benchmark. Finally, several papers have linked accessibility with gene expression using a single large window encompassing upstream regions, the TSS and/or the gene body [11, 20], sometimes combined with a distance-dependent weighting scheme [20]. We included these approaches as a baseline, given that we would expect that any other approach (both CRE-centric or ChromatinHD) should at the very least be able to isolate the predictive from the non-predictive regions within this larger window. As the reference baseline, to which all other models are compared, we included the total number of counts within the full window around the TSS.

#### Comparing motif enrichment

We scanned for motifs within gene regions using position weight matrices and precalculated thresholds from HOCOMOCO v11 [75]. To calculate enrichment of motifs within DARs of a particular cell type, we compared the motif counts from DARs from cell type A with all DARs from other cell types. In this way, we automatically controlled for GC content and bias towards particular regions (gene body, promoter), as these are likely equally present between DARs from different cell types. A Fisher exact test was subsequently used to determine motif enrichment and p-values, which were corrected using the Benjamini-Hochberg procedure. To compare different methods according to their motif enrichment, we extracted the fold-change of differential gene expression using scanpy’s ‘rank_genes_groups’ and calculated correlation between the log-odds-ratio of motif enrichment of a TF with the fold-change of the TF’s gene. To compare ChromatinHD with any CRE-centric DAR method, we defined ChromatinHD DARs as those regions for which log *p*(*x, g*|ct) *>* cutoff_ct_ where cutoff_ct_ was defined such that the same number of positions were deemed differential by both *ChromatinHD-diff* and the method in the particular cell type.

#### Comparing eQTL and GWAS SNP enrichment

We obtained GWAS SNPs linked to immunological disorders (Table S1) from the GWAS catalog [76]. For eQTL enrichment, we obtained eQTLs for whole blood from GTEX [62]. For both GWAS SNPs and eQTLs, we added variants in linkage disequilibrium *r*^2^ ≥ 0.9 using Ensembl’s REST API [77] using the GBR population from phase 3 of the 1000 genomes project [78]. To compare methods according to their eQTL/SNP enrichment, we then calculated the percentage of genes for which at least one variant was present within a DAR. This approach was used to ensure that highly polymorphic genes, e.g. in the HLA locus, were weighted equally as genes with less variation. The direct comparison with ChromatinHD DARs was performed with the same approach as used for benchmarking with motif enrichment.

### 5.8 Overlap between CRE and ChromatinHD methods

We calculated the overlap between differential accessibility in two ways. The “Jaccard positions” score was calculated using the Jaccard similarity *J* between whether a position was part of a differential region. The “F1 regions” score [79] between method *a* and method *b* was calculated by first determining the Jaccard similarity between all pairs of DARs *J*_*g,i,j*_, and then calculating:

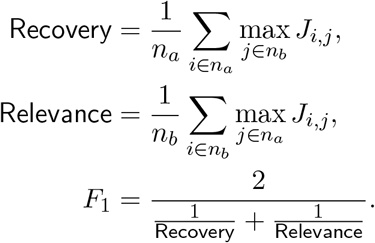

Here, *n*_*a*_ and *n*_*b*_ is defined as the number of DARs in method *a* and *b* respectively.

### 5.9 Co-predictivity

To compare a model where information can be shared between fragments, we first trained a baseline model in which only one linear layer is used to predict gene expression from the cellxgene embedding. After training, we extended the model with an additional gene-specific linear layer with gene-specific parameters with 10 dimensions both as input and output, followed by a ReLU and another linear layer, with gene-specific parameters converting the 10 dimensions into a single scalar value representing gene expression. We summed the output from this layer with those coming from the original linear layer. This model was then re-trained in the same way described above.

To determine co-predictivity between pairs of windows, we used the censoring approach to calculate the predictivity of a particular window in a gene on a per-cell basis ∆*cor*_*c,g,w*_. Specifically, we first z-scored the actual gene expression ***y***_*norm,g*_, predicted gene expression 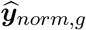 and perturbed gene expression for a particular window 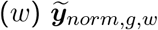 as:

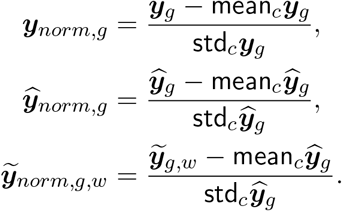

We then calculated the cell-specific predictivity for a window *w* as:

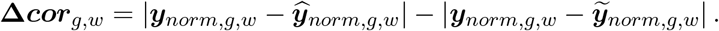

Co-predictivity was then determined by calculating the Pearson correlation between all pairs of windows:

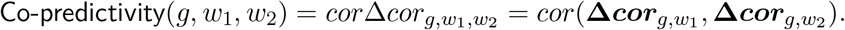

To determine the average co-predictivity for different positions relative to the TSS or different distances, we binned the distances in 2kb bins and calculated the average co-predictivity per bin across all genes. To determine the enrichment of GWAS SNPs in high predictivity (∆*cor*) or co-predictivity (*cor*∆*cor*), we calculated the average absolute co-predictivity values for each gene. Next, for each gene, we calculated the number of SNPs that were present within windows having a (co-)predictivity higher than the *q*-th quantile of (co-)predictivity. We used this to create a global contingency table by summing the enrichment, which ensures that all genes were treated equally regardless of their SNP density. We used this contingency table to calculate the odds-ratio. We used the same set of GWAS SNPs as was used for the benchmark section.

### 5.10 Linking DNA contact with co-predicitivity

We obtained Hi-C contact matrices for Rao 2014 [27] from https://data.4dnucleome.org/files-processed/4DNFIXP4QG5B/ and for Harris et al. 2023 [41] from https://www.encodeproject.org/files/ENCFF555ISR. For the Rao et al. 2014 data, we mapped each 1kb bin to co-predictivity windows (100 bp) based on the largest overlap in individual positions, and subsequently extracted the maximal absolute co-predictivity value for each 1kb bin. For the Harris et al. 2023 data, we applied a similar procedure but using 500 bp bins.

To determine the correspondence between Hi-C data and co-predictivity, we first stratified all pairs of regions in different distance bins of size 10kb up to a maximum of 150kb, which corrected the analysis both for the distance-dependency of both Hi-C scores and co-predictivity (**Figure 4f**). Per gene and per distance bin, we then calculated an odds score based on whether co-predictivity and/or the normalized Hi-C contact score were higher than their respective mean scores. Log odds-ratios were then averaged across genes and distance bins. Max-pooling on the Hi-C data was performed by taking the maximal Hi-C contact score for all region pairs within a specified manhattan distance.

We calculated pile-ups of Hi-C signal around co-predictive regions by first selecting only those pairs with *cor*∆*cor >* 0 and within a specific distance bin, i.e. 5-10kb, 10-15kb, 20-25kb, 30-35kb and 40-45kb. For each such pair, we picked a random pair of regions with the same genomic distance within the gene’s window (i.e. -100kb to +100kb), and calculated the ratio between the co-predictive pair and random pair. These log-ratios were averaged to retrieve the final pile-ups as in **Figure 4i**. Individual examples, as shown in **Figure ED4e**, were created by normalizing against 100 random pairs from the same gene. To determine the change in pileups between two sets of genes, we calculated the change in log-ratio between co-predictive regions coming from genes that were significantly upregulated in B-cells (naive, memory and plasma cells, fold-change ≥ 2) to all other genes.

To determine the enrichment of looping factors (YY1, CTCF, RAD21, ZNF143), B-cell TFs (SPI1, EBF1, IRF4, PAX5) or the transcription machinery (POLR2A), we obtained ChIP-seq data from GM12878 cells from ENCODE [9] (Table S2). We ranked all 100 bp regions for a gene according to their predictivity (∆*cor*) subsetting on those with ∆*cor* ≤ −0.01, and iteratively selected those regions if they were further away than 1kb from any previously selected region. With these regions as the center, we calculated the mean log-ratio between highly co-predictive enhancers (those where average co-predictivity was bigger than the mean within a gene) versus lowly co-predictive enhancers (all other enhancers) for various up and down-stream distances.

### 5.11 Fragment sizes

To compare the enrichment of motifs depending on predictivity or Mono−/TF footprint ratios, we calculated the GC-content corrected motif counts by first creating a spline model (quadratic spline, 8 knots, implemented in R’s ‘smooth.spline’ function) that calculates the relationship between motif counts and GC content in 100 bp windows. From this model, we extracted the residuals to get the corrected counts. Next, we used linear models (using R’s ‘lm’ function) for each motif to determine the relationship between either predictivity or Mono−/TF footprint ratio with these GC-corrected counts. The slope and its associated p-value (corrected using Benjamini-Hochberg correction) were used for downstream analysis.

For footprinting, we used Hint-ATAC [54], using the Regulatory Analysis Toolbox (RGT) toolkit v1.0.0 (https://reg-gen.readthedocs.io/en/latest/), in ATAC-seq and paired-end mode. We obtained footprints using DNase I hypersensitivity footprinting from different immune cell types from [80].

As reference ChIP-seq data, we downloaded all ChIP-seq peaks from ENCODE using the GM12878 cell line and excluding CTCF, RAD21, POL2RA and EP300 (Table S2). To count the number of TFs binding within a window, we determined if at least part of a peak, as identified by ENCODE’s narrowPeaks output, overlapped with the window.

## 6. Data availability

Multiome data was obtained from https://www.10xgenomics.com/resources/datasets. ENCODE ChIP-seq data from https://www.encodeproject.org/ (Table S2) [9]. DNase I footprinting data from https://zenodo.org/record/3905306 [80]. Hi-C data from https://data.4dnucleome.org/files-processed/4DNFIXP4QG5B [27] and https://www.encodeproject.org/files/ENCFF555ISR [41].

## 7. Code availability

The ChromatinHD python package is available at https://github.com/DeplanckeLab/ChromatinHD and will be published on pip and bioconda. The pipeline to reproduce the main results from the manuscript will be published at https://github.com/DeplanckeLab/ChromatinHD_manuscript.

## 8. Extended data figures

**Figure ED1:**
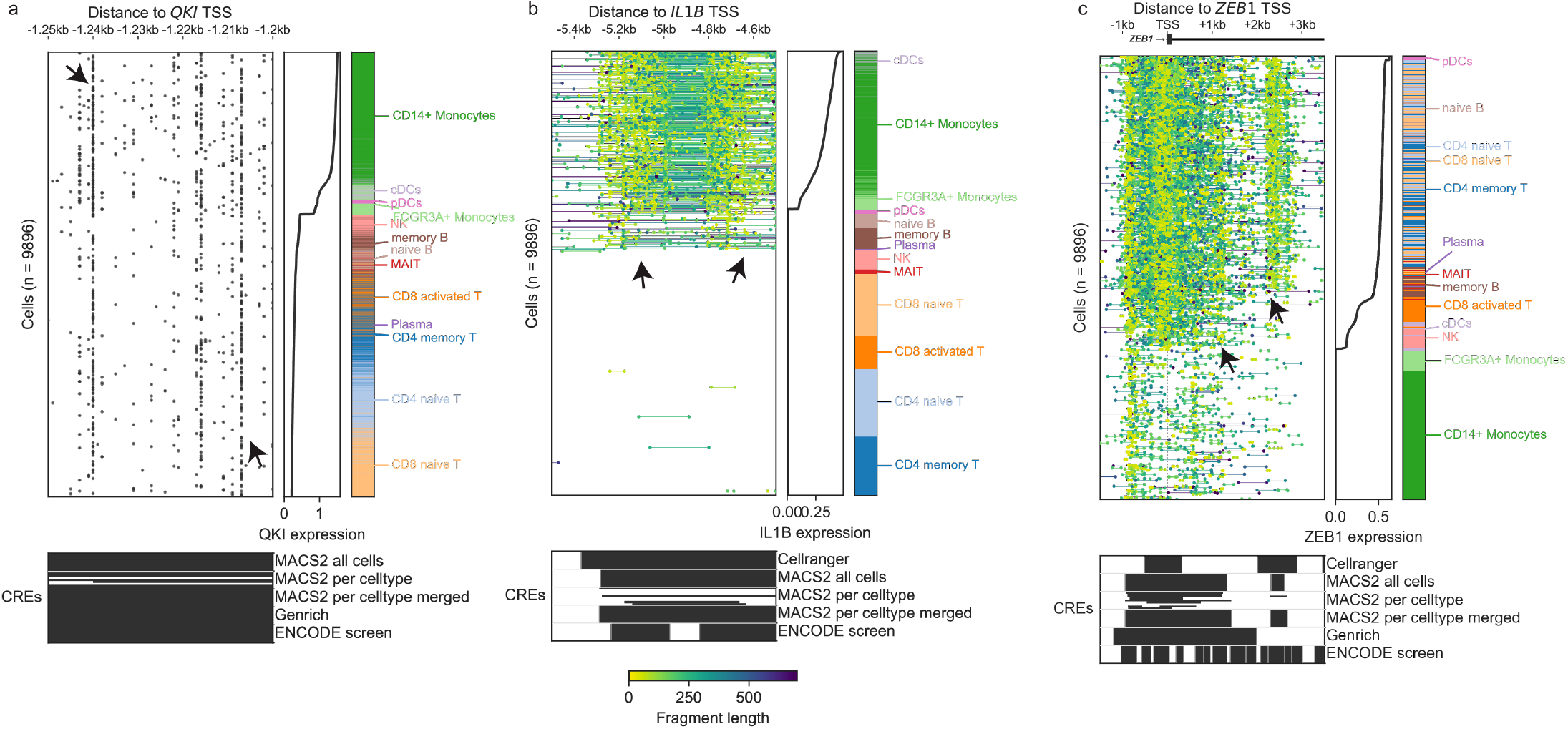
Examples of chromatin accessibility changes at 3 different resolutions. (**a**) A narrow region of 50 base pairs upstream of the *QKI* transcription start site (TSS), where each dot represents a cut site. Between cell types with low or high expression, the accessibility shifts from -1.21kb towards -1.24kb. (**b**) A region of 1kb upstream of the *IL1B* TSS, with individual fragments connected by a line and fragment sizes indicated by hue. Two regions of high expression-specific accessibility can be observed, and fragments often connect these two regions along a 200bp region without cut sites, likely a resident nucleosome. Note that the two regions are slightly different in size, and how peak-based methods combine both regions in a common peak despite the right region being more accessible in B-cells. (**c**) A region of 5kb around the *ZEB1* TSS, with extensive changes both up and down-stream of the TSS, some of which being specific to low and high expression, but often grouped together into common cis-regulatory regions (CREs). The arrows indicate respective regions of interest.

**Figure ED2:**
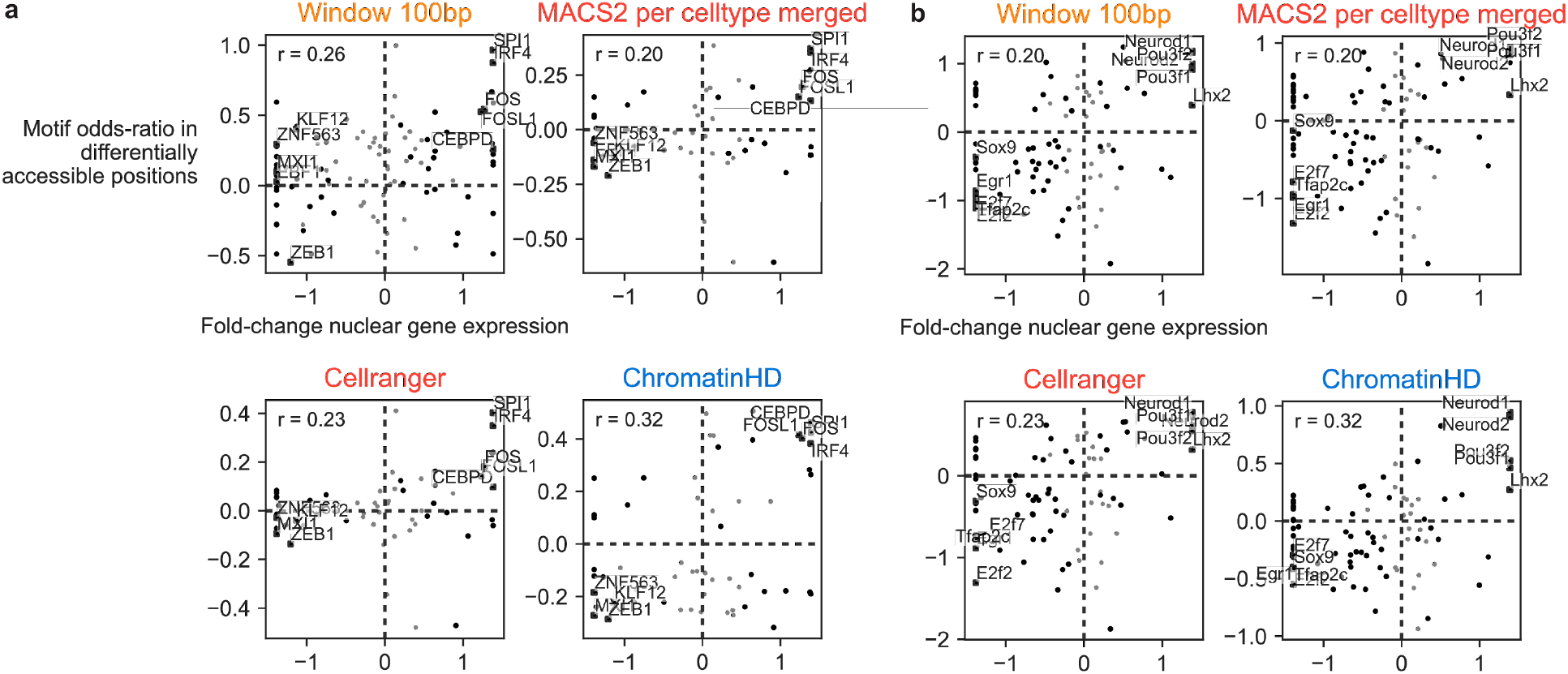
Relationship between differential gene expression and motif enrichment in differentially accessible regions. (**a**) In conventional dendritic cells from the pbmc10k dataset. (**b**) In neural progenitor cells in the e18 mouse brain dataset.

**Figure ED3:**
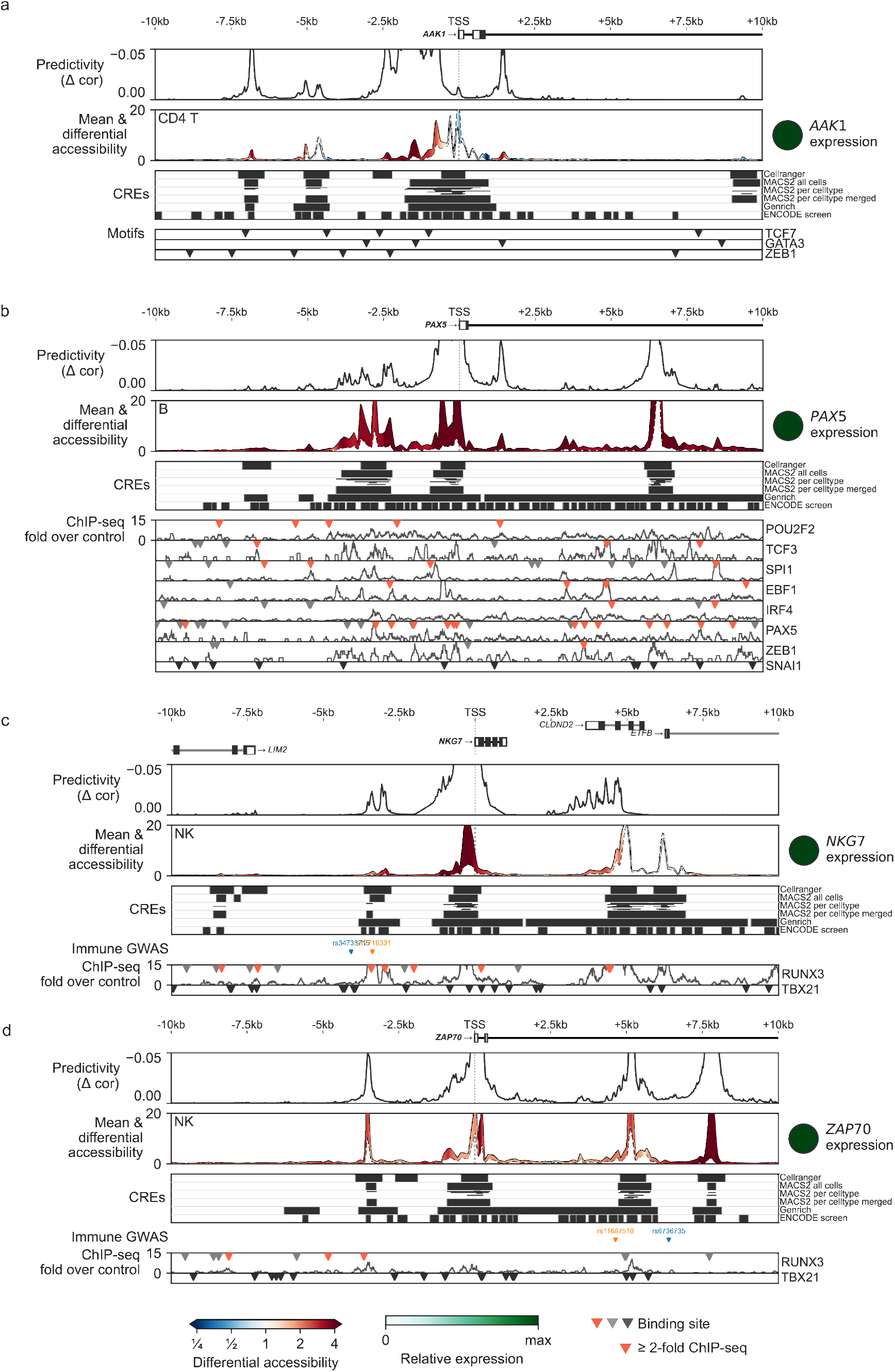
Additional examples of ChromatinHD model interpretation. Positional interpretation of both *ChromatinHD-pred* (Predictivity, top) and *ChromatinHD-diff* (Mean and differential accessibility, middle) across different genes, along with positions of immune-related GWAS SNPs (colors representing different haplotypes with LD relevant transcription factors and ChIP-seq tracks if available.

**Figure ED4:**
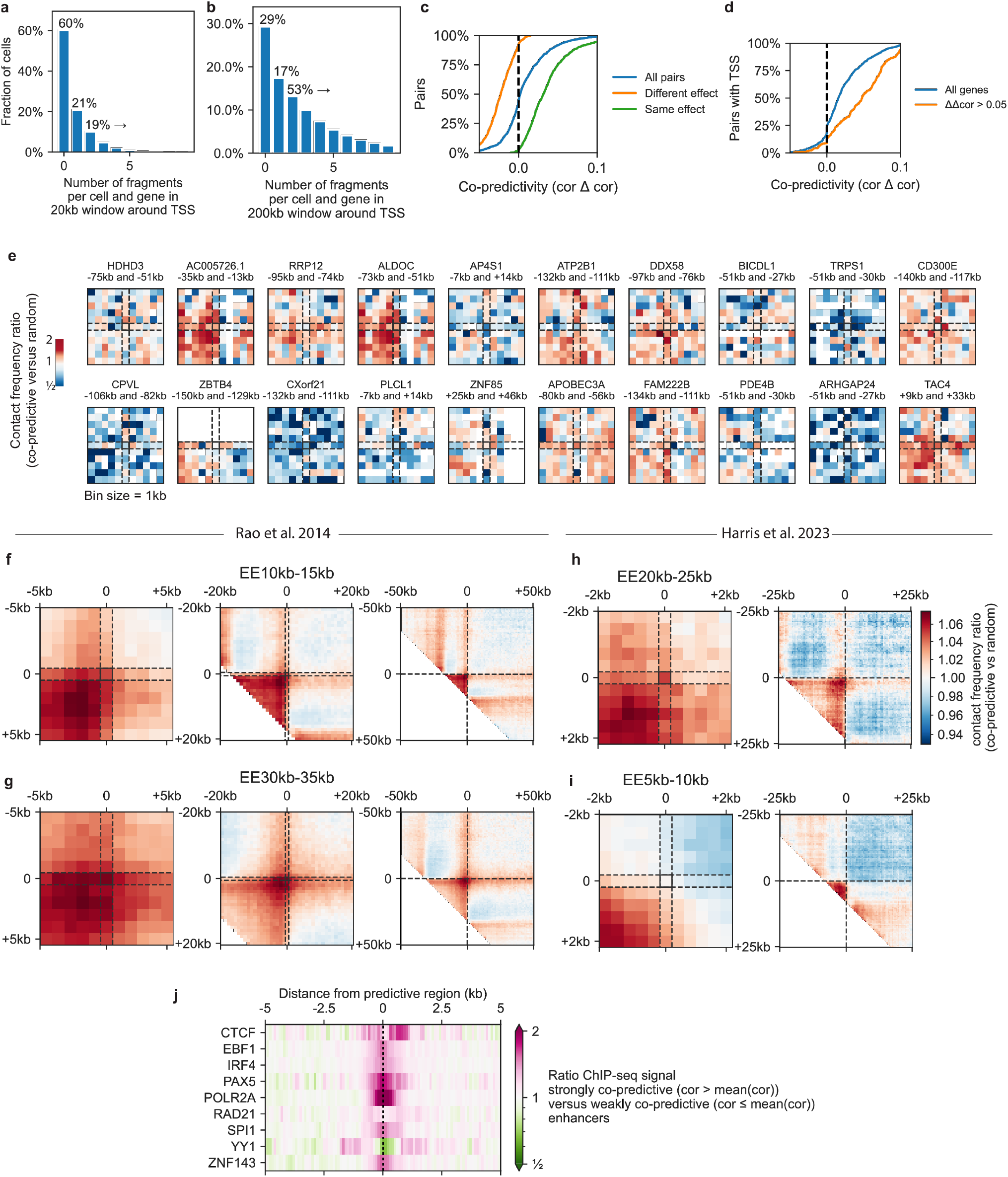
Statistics on co-predictive regions. (**a**) Distribution of fragments located within the -10kb and +10kb region from the transcription start site (TSS). (**b**) Same as *a* but for -100kb and +100kb regions. (**c**) Distribution of co-predictivity between windows that have the same effect (green, as determined by *ChromatinHD-pred*) or opposite effect (orange). (**d**) Distribution of co-predictivity based on the difference in predictive accuracy in the additive versus non-additive model from **Figure 4a**. (**e**) Hi-C results between the strongest co-predictive regions for several genes in the pbm10k dataset. Data is normalized for random pairs with the same genomic distance close to the same gene. (**f-i**) Hi-C pile-ups of potential DNA contact points (C1 and C2) close to two co-predictive regions (E1 and E2, cor ∆ cor > 0). Shown is the relative Hi-C signal centered on the co-predictive pair divided by a random pair around the same gene with the same genomic distance. (**f**) Pile-up where the distance between the co-predictive pair (E1-E2) is between 10 and 15 kb. Bin size 1kb. (**g**) E1-E2 distance between 30 and 35kb (**h**) Pile-up for [41] with E1-E2 distance between 20 and 25 kb. Bin size 500 bp. (**i**) E1-E2 distance between 5 and 10kb. (**j**) Enrichment of binding of different cell type-specific transcription factors and looping factors at or close to predictive regions. ChIP-seq data from GM12878, a B-cell-like cell line [13]. Shown is the (log-)average ratio of signal-to-background.

**Figure ED5:**
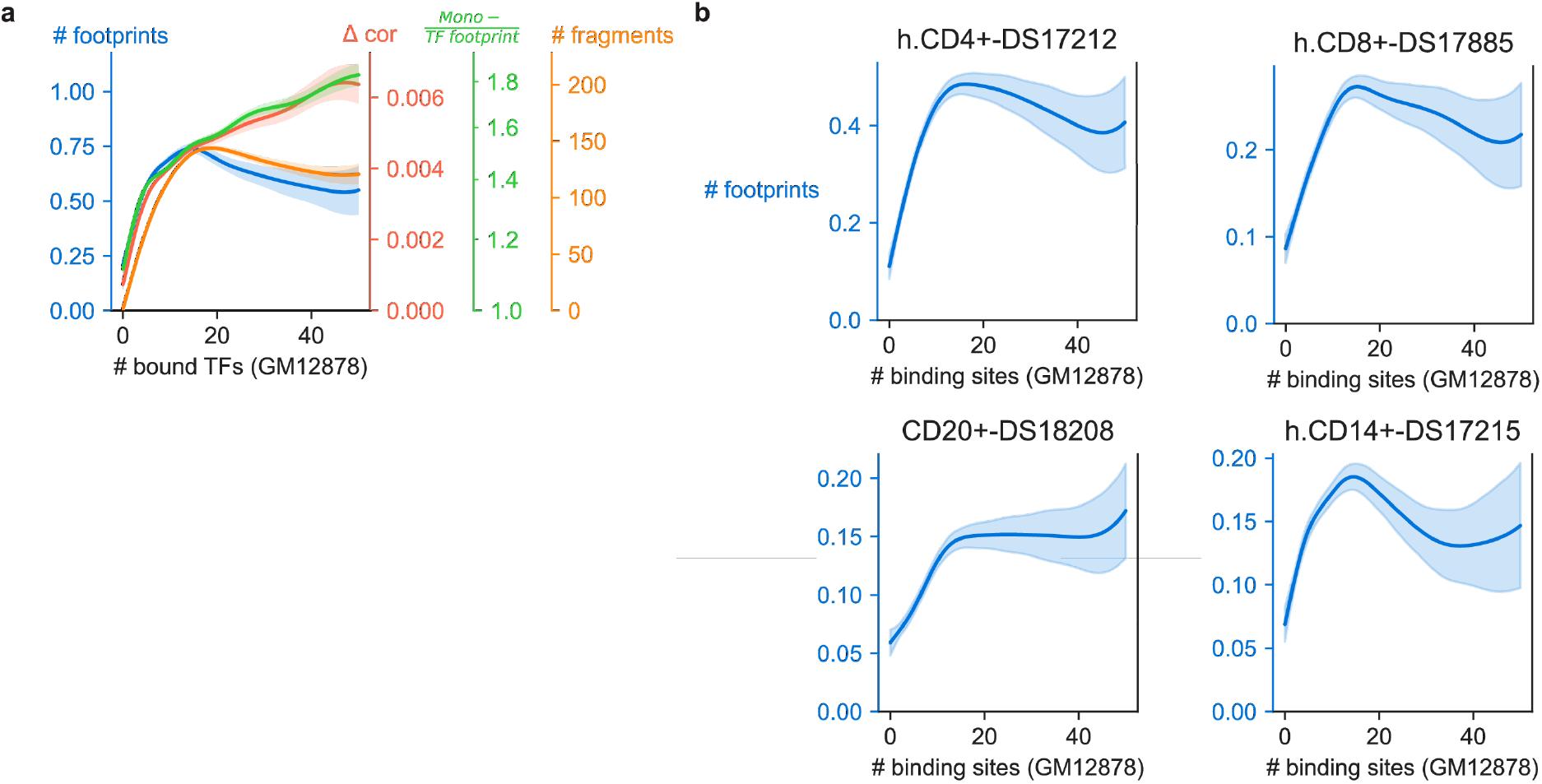
Comparing the density of transcription factor binding with the number of detected footprints, predictivity and number of fragments. (**a**) Relationship of the # of (indirectly) bound transcription factors within a 100 bp window according to ENCODE GM12878 data and the # of footprints according to HINT-ATAC on the pbmc10k data (blue), predictivity (red), ratio of Mono− versus transcription factor footprint fragments (green) and overall number of fragments (orange). Shown is the mean and standard error of a spline fit using R’s gam function with smoothing parameter *sp* = 1. ChIP-seq data of all transcription factors (n=132) were used, excluding looping factors (CTCF, RAD21, YY1). (**b**) Relationship between the density of transcription factor binding (x-axis) in ENCODE GM12878 data, and the number of footprints detected in DNase I hypersensitivity data [55] in various cell types.

## 9. Supplementary figures

**Figure S1:**
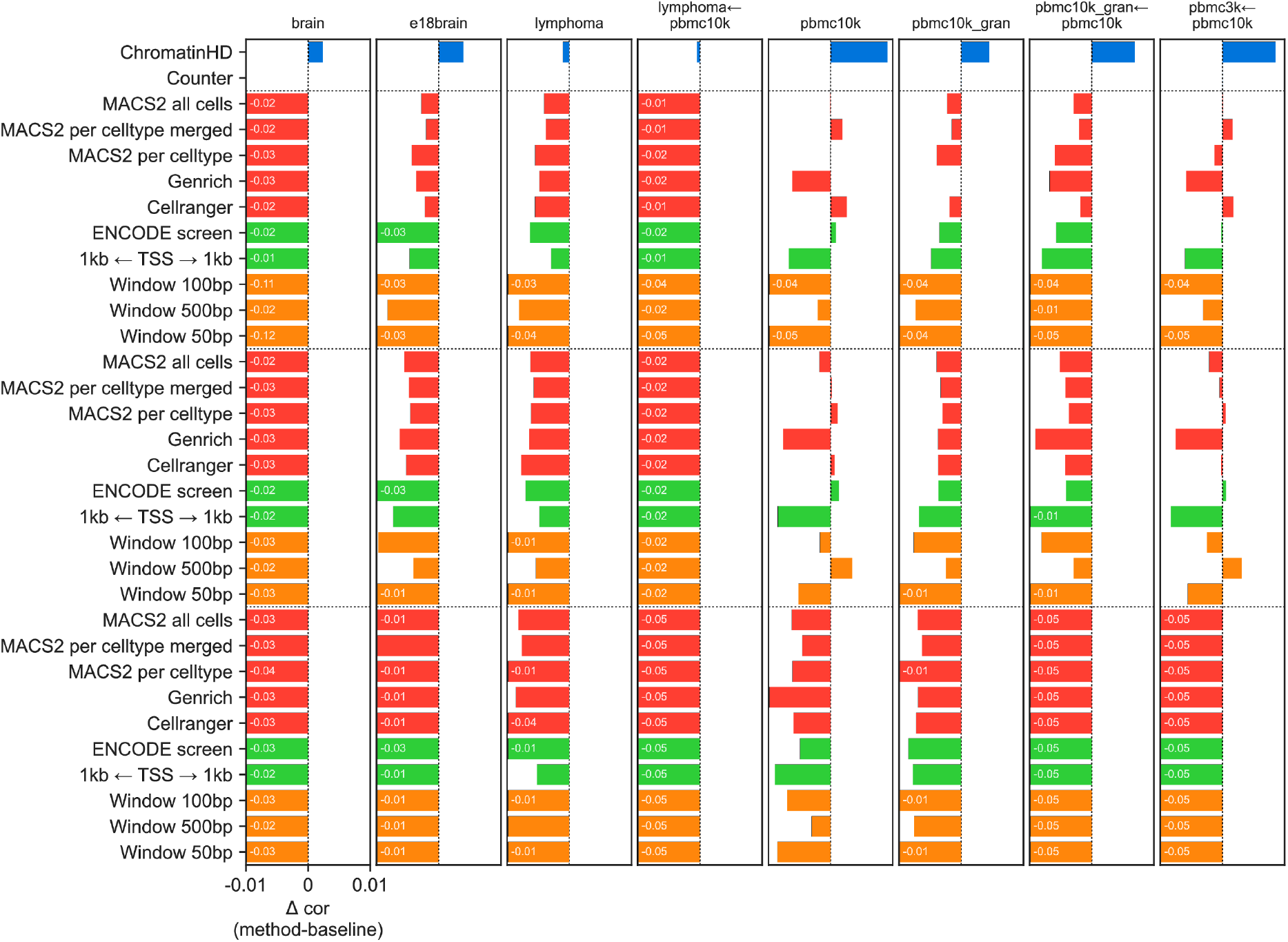
Comparing the predictive accuracy on test data of *ChromatinHD-pred*. Test data either includes cells from the same dataset (if only one dataset is shown at the top) or cells form another dataset (test dataset ← train dataset).

**Figure S2:**
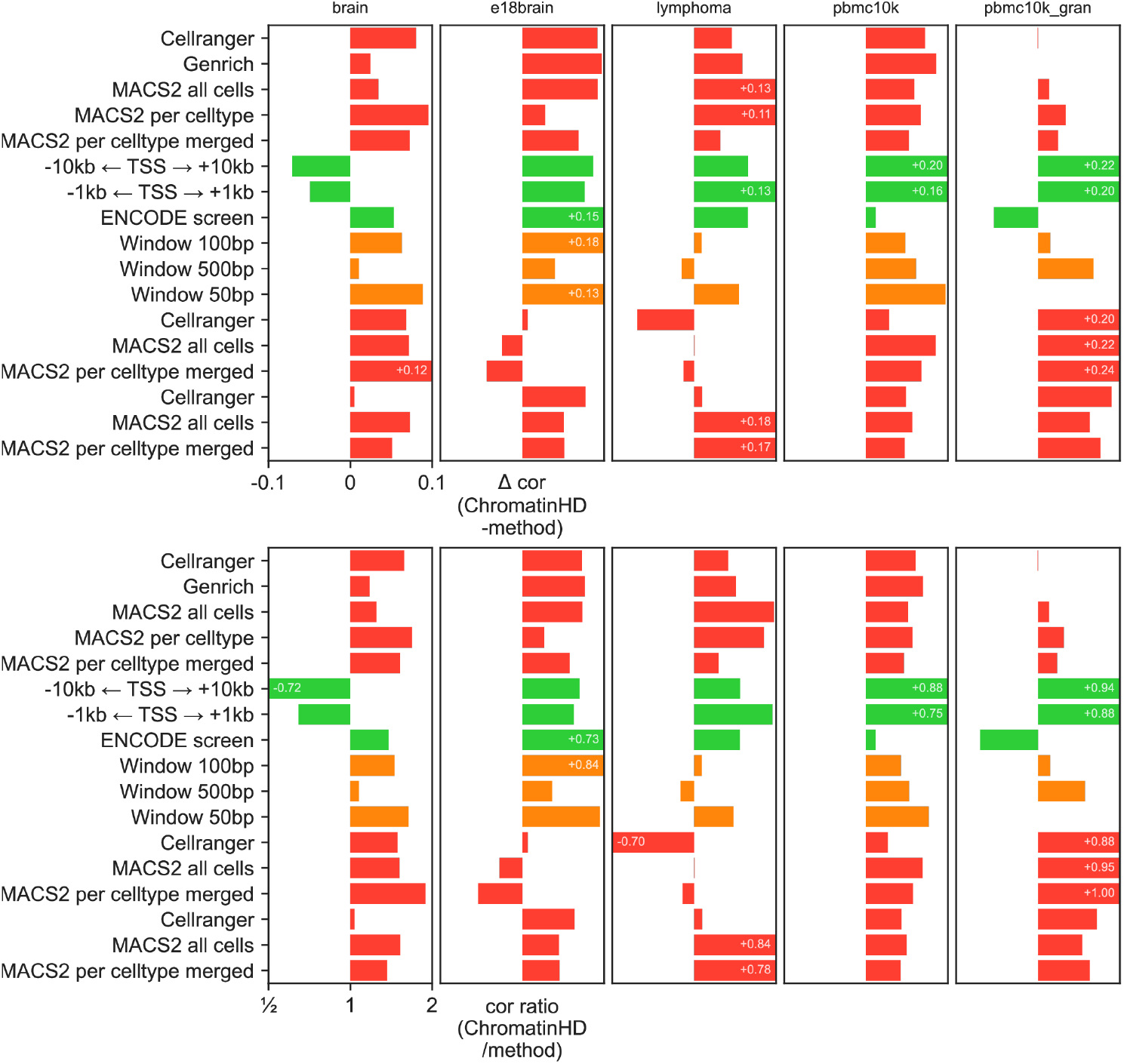
Comparing the correspondence between motif enrichment and differential expression for *ChromatinHD-diff*compared to other differential accessibility methods.

**Figure S3:**
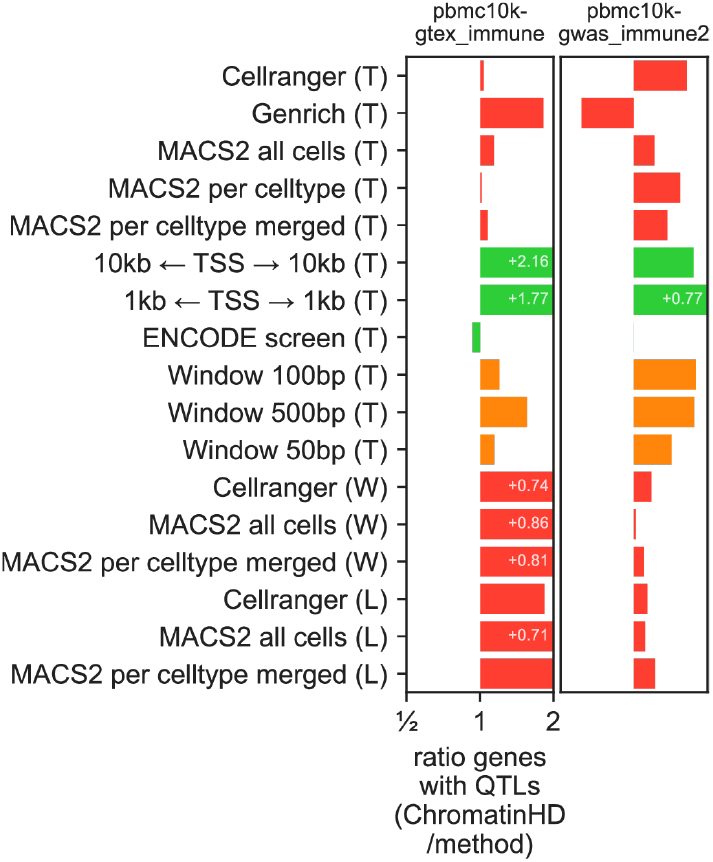
Comparing the percentage of genes for which a known immune-related GWAS SNP or eQTL is located within a differentially accessible region.

**Figure S4:**
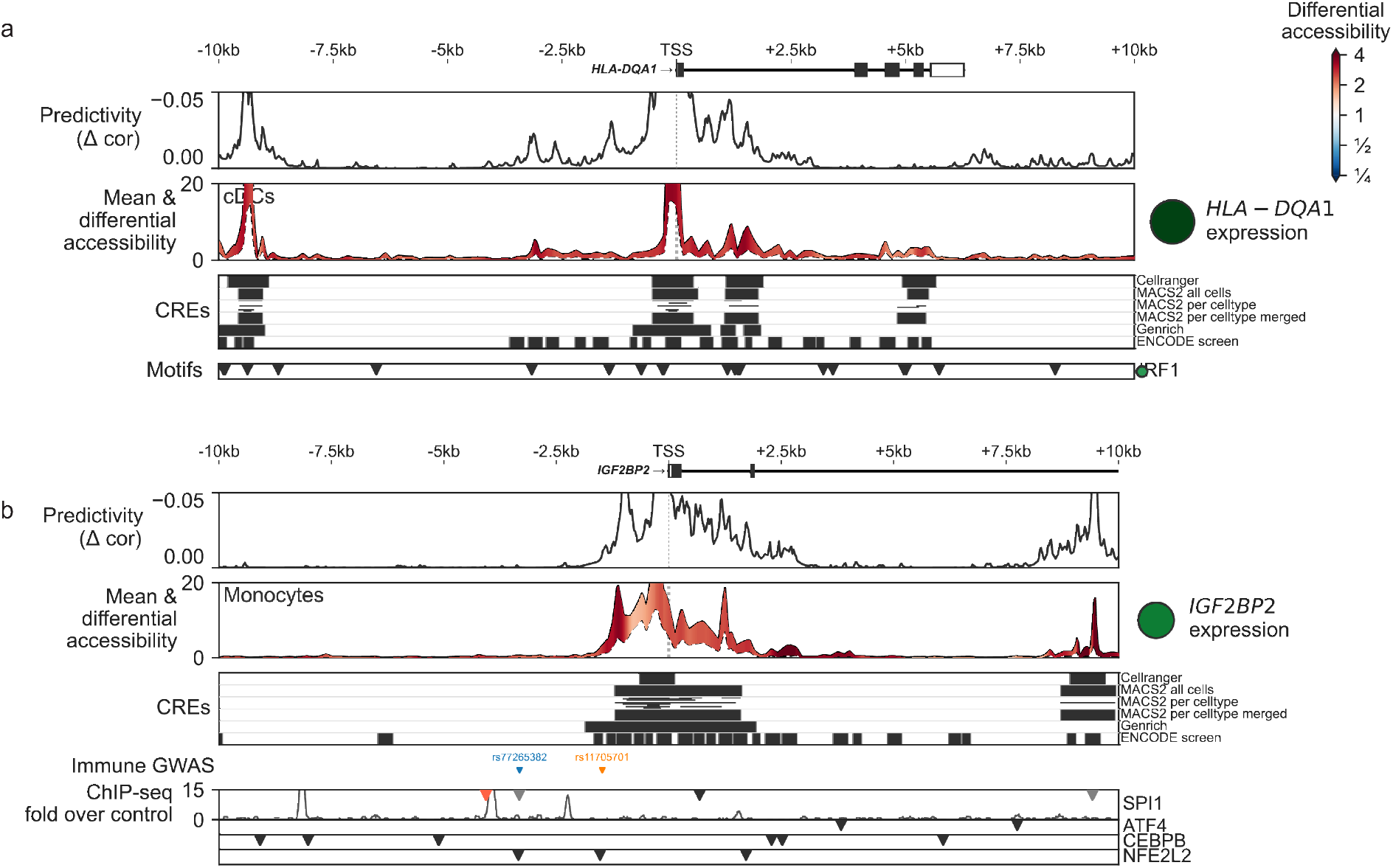
Additional examples of ChromatinHD model interpretation. Positional interpretation of both *ChromatinHD-pred* (Predictivity, top) and *ChromatinHD-diff* (Mean and differential accessibility, middle) models across different genes, along with positions of immune-related GWAS SNPs (colors representing different haplotypes with LD transcription factors and ChIP-seq tracks if available.

## 11 Additional information

## Acknowledgements

This work was supported by the European Union’s Horizon 2020 research and innovation program under the Marie Skłodowska-Curie grant agreement 101028476 (to W.S.), as well as by SNSF Project Grant #310030_197082 funding (to B.D.). We thank Vincent Gardeux, Judith Kribelbauer and Guido van Mierlo for their helpful feedback on the manuscript.

## Author contributions

W.S. and B.D. conceived and designed the study. W.S. and O.P. analyzed the data. W.S. and B.D. wrote the manuscript.

