## Supplementary Note 1 for "ChromatinHD connects single-cell DNA accessibility and conformation to gene expression through scale-adaptive machine learning"

### Supplementary note 1: Overall comparison of predictive or differential regions identified by ChromatinHD versus CRE-centric approaches.

We found relatively little overlap between ChromatinHD and CRE-centric methods in terms of differentially accessibly individual base pairs and regions (20-30% depending on the dataset), lower than the overlap between differential peaks called by different peak-calling approaches (20%-60%) (**Figure 1.1a**). The difference between the two approaches was particularly striking around the TSS where the overlap with differential MACS2 peaks fell below 20% (**Figure 1.1b**). MACS2 peaks had a clear negative bias towards the -2.5kb and +2.5kb regions around a TSS, while ChromatinHD found DARs with a nearly monotonic decrease starting from the TSS in both up- and downstream direction (**Figure 1.1b**). This difference between differential peaks and ChromatinHD was present across datasets, cell types, peak callers and differential accessibility algorithms (**Figure 1.1c**), indicating that this is caused by some common bias among CRE-centric approaches. The apparent bias was also present, albeit to a lesser extent, in sliding window and pre-defined regions (**Figure 1.1c**).

We posited that the first explanation for the discrepancy between differentially accessible peaks and ChromatinHD DARs could derive from the definition of a background. Usually, peaks are identified as areas that stand out from the background signal, although the exact definition of this background depends on the peak caller that is used [1–3]. We hypothesized that this background assumption may negatively impact moderately accessible regions that are near highly accessible ones, a frequent occurrence around the promoter, even though those moderately accessible regions might well be functional. Indeed, using ChromatinHD, we found numerous examples of regions that clearly change in accessibility in a particular cell type, but that were not called as differential (**Figure 1.2a**). These regions often appear adjacent to one or more existing regions with higher accessibility (**Figure 1.2a**), which likely caused them to be incorrectly dismissed as background.

To test whether this is true in general, we used ChromatinHD's differential density distribution to quantify the mean and variation in accessibility for each position. We found that the maximal accessibility of differential peaks is nearly twice as high as that of differential ChromatinHD regions (**Figure 1.2b**), both at and outside of the TSS (**Figure 1.2c**), while there was no remarkable difference in average accessibility (**Figure 1.2b**). We hypothesized that, given that highly accessible regions are open in many cell types/states, they may be less likely to confer specificity and are therefore less likely to be directly involved in cell type-specific TF binding and gene regulation. To test this, we used ChromatinHD's differential accessibility distribution to bin individual positions according to their mean and differential accessibility. We found that globally, the positions with intermediate accessibility are the most variable, with a clear drop in differential accessibility when positions are either highly or lowly accessible on average (**Figure 1.2d**). This disconnect between mean accessibility and differential accessibility can be clearly seen in individual examples, with for example *BCL2* and *QKI* only showing minimal changes in their most accessible regions (often around the TSS) (Figure 3). Yet these genes do exhibit strong changes in gene expression and, interestingly, also accessibility changes in moderately accessible regions. To test whether these regions would indeed confer cell type-specificity, we did motif enrichment of cell type-specific TFs for different combinations of mean and differential accessibility. We found that binding sites of cell type-specific TFs are equally, and sometimes more, enriched in regions of moderate accessibility even when stratified according to the magnitude of differential accessibility (**Figure 1.2e-f**). This was mostly consistent across TF-cell type combinations, albeit with some exceptions such as *TBX21* in NK cells and *NFKB2* in B-cells (**Figure 1.2f**). Altogether, this indicates that moderately accessible regions may be as, if not more relevant, than highly accessible ones in mediating cell type-specific regulation. This is in contradiction with the use of background to distinguish functional from non-functional accessible regions.

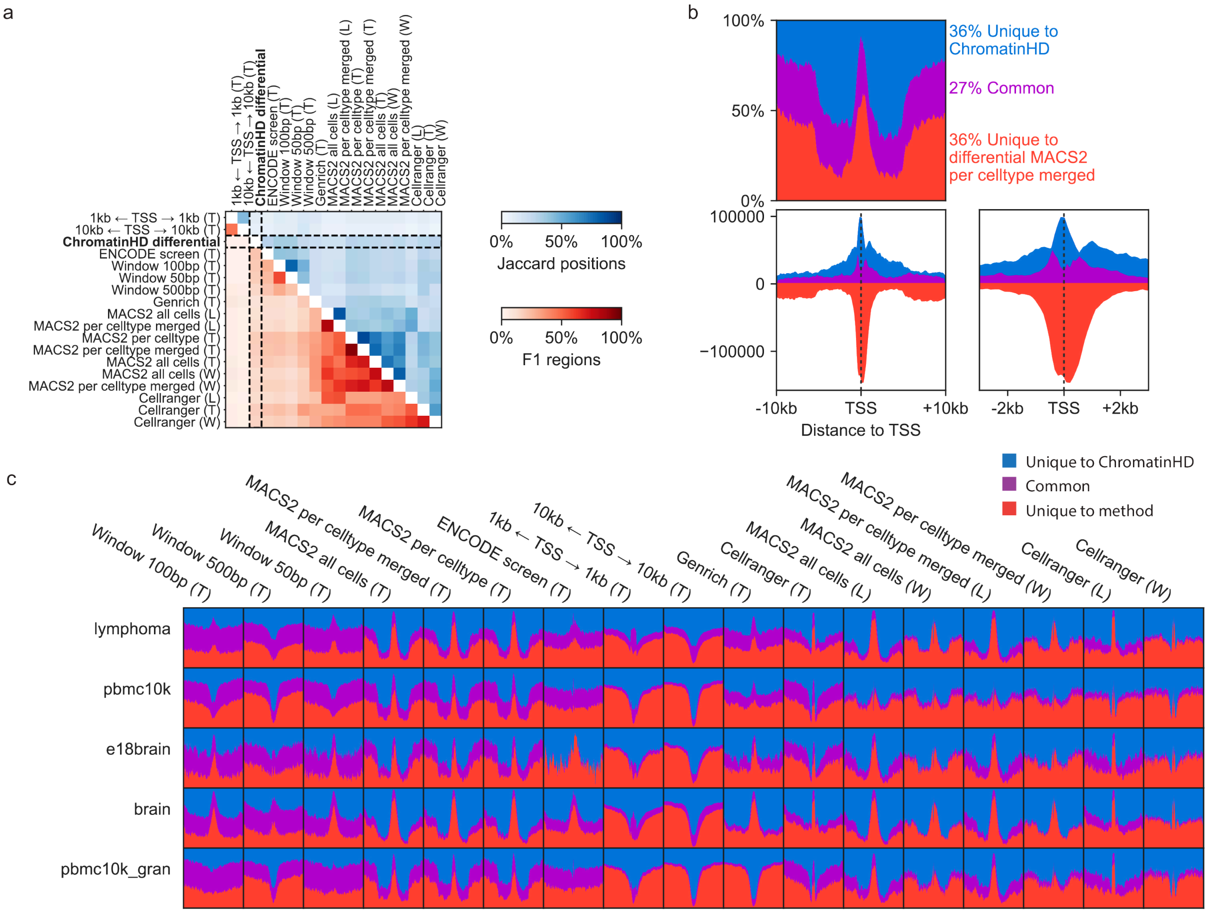


***Figure 1.1:*** **Overlap between ChromatinHD and CRE-centric approaches**. We compare with "MACS2 per celltype merged" and the pbmc10k dataset if not specified otherwise. (**a**) Overlap between various peak-calling + differential accessibility methods. The upper triangle shows the Jaccard index on absolute positions, while the lower triangle shows the overlap as an F1 score between regions. (**b**) Relative (up) and absolute (down) overlap between differential ChromatinHD and differential accessible peak positions across all genes in the pbmc10k dataset. Bin size = 100bp. (**c**) Relative overlap between *ChromatinHD-diff* regions and differentially accessible peak positions across all genes in various peak callers (top) and datasets (left). Bin size = 100bp.

The background assumption can also affect predictive models, as regions predictive for gene expression may be dismissed as background. When comparing the predictivity of individual 100 bp windows, about 27% of predictivity is contributed by regions outside any peak, although this percentage varies considerably between genes (**Figure 1.2g**). If we considered the top 10% of predictive positions across all genes, 49.4% of positions were not located within a peak, a percentage that is consistent even when considering only fragments upstream of the gene body (**Figure 1.2h**). Furthermore, the correlation between overall accessibility and predictivity was highly variable across genes, with over half of the genes having a correlation lower than 0.5 (**Figure 1.2i**). This was particularly low for genes where ChromatinHD predicts gene expression much better than the baseline (**Figure 1.2i**, orange). Altogether, this highlights that mean accessibility is not necessarily an optimal indicator for predicting the impact of a region on gene expression, which violates the signal-over-background assumption made by peak-calling methods.

A second reason for the difference between CRE-centric approaches and ChromatinHD is that CREs may carry a substantial number of positions that are neither differential nor predictive. These "bystander" positions would reduce statistical power, increase noise, and may not confer specificity as the accessibility is locally not changing between cell types. To quantify this on a per-region basis, we calculated the fraction of positions within a differential peak that was called differential by ChromatinHD. Overall, we found that the percentage of non-differential positions was very variable, with 18.6% of the differential MACS2 peaks containing no differential positions, while 32.0% was fully differential (**Figure 1.3a**). A majority of differential peaks, 58.9%, contained less than 50% differential positions (**Figure 1.3a**), a statistic particularly high in larger peaks (>1kb). The latter often encompassed regions with broad overall accessibility, of which only a small part truly changed in accessibility (**Figure 1.3b**). While this bias was present both in proximal and distal positions from the TSS, the number of bystander positions was substantially stronger close to the TSS (**Figure 1.3c**). Window-based approaches equally suffered from an increase in bystander positions (**Figure 1.3d**), given that these methods also group all positions within a window together. Similarly, bystander positions also impact predictive models. For 48.5% of the most predictive peaks (MACS2 cell type merged peaks), the summit of the peak did not match the most predictive position according to ChromatinHD within a 100 bp window. In fact, for about 11% of peaks the most predictive region was more than 500bp from the summit (**Figure 1.3e**). Overall, this indicates that there is substantial heterogeneity within CREs, and that aggregating information over a CRE, even when centered at the summit, removes important gene regulatory information.

Finally, we note that the overlap between CRE-centric methods is low (<50%) (**Figure 1.1a**). This indicates that peak-calling methods themselves are already in disagreement on how CREs should be defined, i.e. what regions should constitute a background, and how accessibility regions should be separated into different peaks.

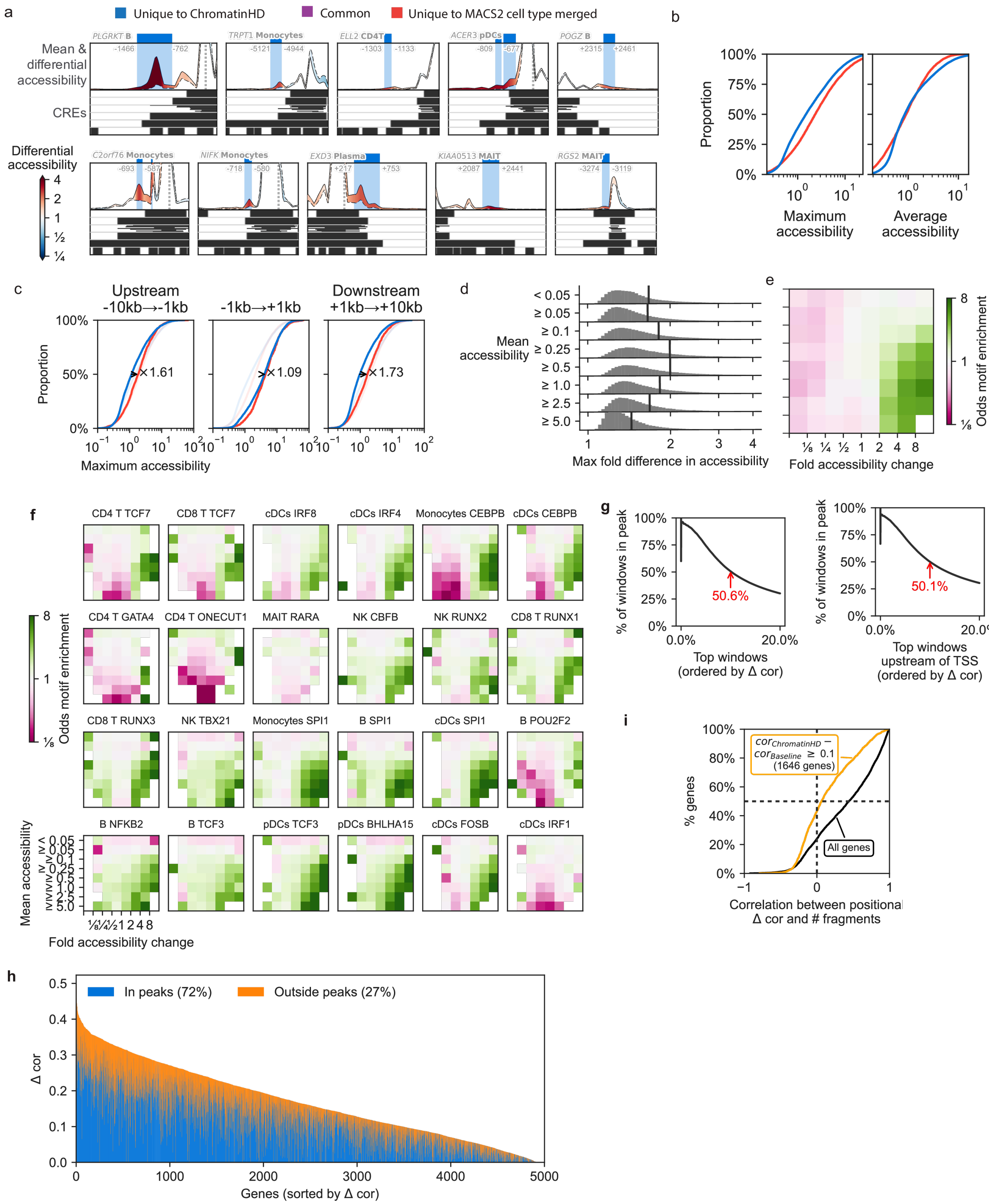


***Figure 1.2:*** Caption next page

***Figure 1.2:*** **Exploring the background bias**. (**a**) Examples of regions that were deemed differentially accessible by ChromatinHD but not by MACS2 per cell type merged even though they are close to another, more accessible, region. The grey line denotes the TSS. (**b**) Distribution of average and maximum accessibility for each differential region. Colors as in *a*. (**c**) Maximum accessibility (as determined by *ChromatinHD-diff*) in *ChromatinHD-diff* regions and differential peaks, in regions upstream, at or downstream of the transcription start site (TSS). (**d**) Distribution and average values (vertical line) of the maximal accessibility fold-difference across cell types/states for each position, binned by the mean accessibility (across cell types/states). (**e**) Average odds-ratio for motif enrichment in combinations of motifs and cell types for pairs of mean and differential accessibility. (**f**) Motif enrichment of cell type-specific transcription factors in different ChromatinHD regions stratified by mean accessibility (y-axis) and differential accessibility (x-axis, fold-change). (**g**) Percentage of top predictive windows (according to *ChromatinHD-pred*, 100 bp) that are contained within a peak, both for windows in the whole gene region (-10kb to +10kb, left), as well as regions upstream of the gene's TSS (-10kb to TSS, right). (**h**) Predictivity ($\Delta cor$) for windows (100 bp) contained within and outside peaks according to *ChromatinHD-pred*. (**i**) Distribution of the correlation between a window's (100 bp) predictivity ($\Delta cor$) and number of fragments. Only windows containing at least 1% of the fragments were considered.

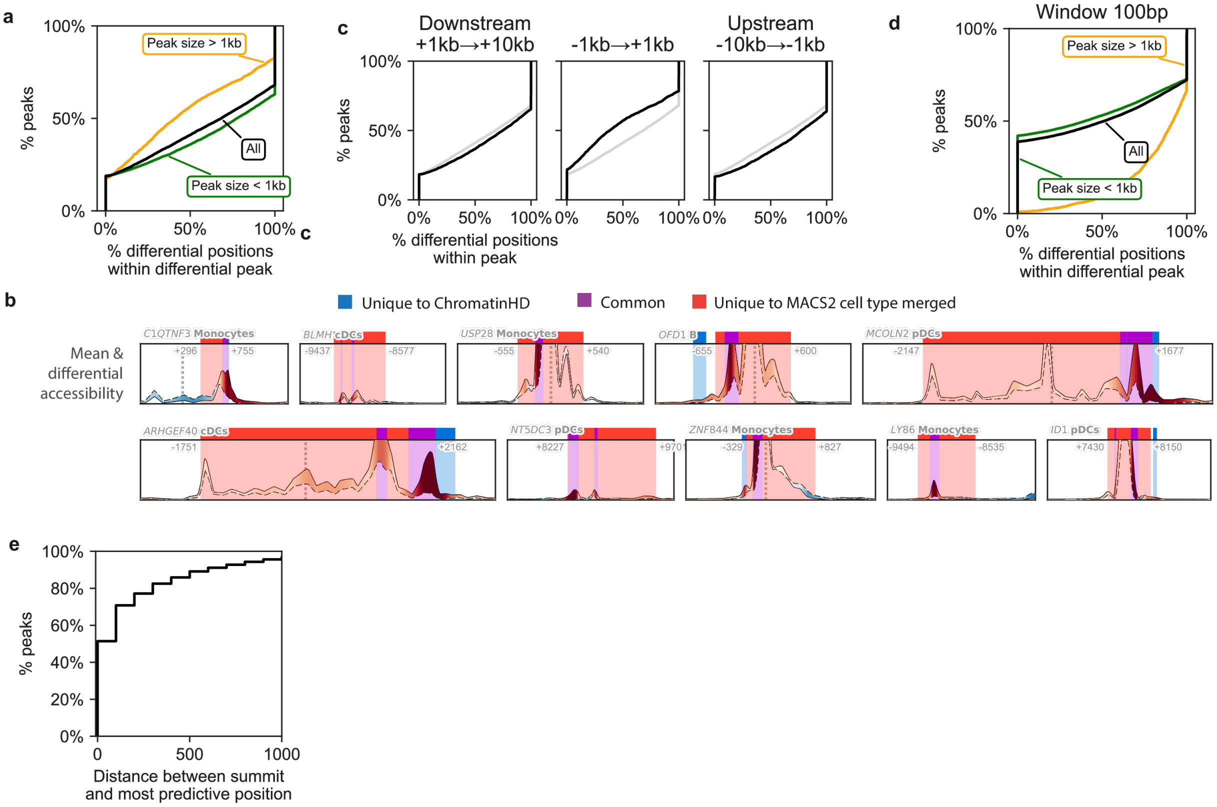


***Figure 1.3:*** **Exploring the bystander bias**. Results from the pbmc10k dataset and t-test MACS2 celltype merged peaks if not specified otherwise. (**a**) Percentage of top predictive windows (according to *ChromatinHD-pred*, 100 bp) that are contained within a peak. (**b**) Examples of differentially accessible peaks (red+purple, at least 1.5x fold-change), of which *ChromatinHD-diff* deemed only a part differentially accessible (at least 1.5x fold-change, blue+purple). The grey line denotes the TSS. (**c**) The number of differential positions (i.e. non-bystander positions), stratified according to relative position to TSS. Differential positions are defined as positions in differential peaks where the fold-change in accessibility $< 2$ according to *ChromatinHD-diff*. (**d**) The number of differential positions as in *a* stratified according to peak size. (**e**) Distance between the most predictive position within a peak (window size 100 bp) and peak summit, defined as the window containing the most cut sites.

3. *Cell Ranger ATAC Algorithms Overview -Software -Single Cell ATAC -Official 10x Genomics Support* https://support.10xgenomics.com/single-cell-atac/software/pipelines/latest/algorithms/overview#peaks (cit. on p. 1).
